# Cue-specific remodeling of the neuronal transcriptome through intron retention programs

**DOI:** 10.1101/2021.10.13.463312

**Authors:** Maxime Mazille, Peter Scheiffele, Oriane Mauger

**Affiliations:** Biozentrum of the University of Basel, Spitalstrasse 41, 4056 Basel, Switzerland

## Abstract

Sub-cellular compartmentalization through the nuclear envelope has for a long time been primarily considered a physical barrier that separates nuclear and cytosolic contents. More recently, nuclear compartmentalization has emerged to harbor key regulatory functions in gene expression. A sizeable proportion of protein-coding mRNAs is more prevalent in the nucleus than in the cytosol reflecting the existence of mechanisms to control mRNA release into the cytosol. However, the biological relevance of the nuclear retention of mRNAs remains unclear. Here, we provide a comprehensive map of the subcellular localization of mRNAs in mature neurons and reveal that transcripts stably retaining introns are broadly targeted for nuclear retention. We systematically probed these transcripts upon neuronal stimulation and found that sub-populations of nuclear-retained transcripts are bi-directionally regulated in response to cues: some appear targeted for degradation while others undergo splicing completion to generate fully mature mRNAs which are exported to the cytosol to increase functional gene expression. Remarkably, different forms of stimulation mobilize distinct groups of intron-retaining transcripts and this selectivity arises from the activation of specific signaling pathways. Overall, our findings uncover cue-specific control of intron retention as a major regulator of acute remodeling of the neuronal transcriptome.

## Introduction

Transcriptome remodeling plays a major role in cellular differentiation and plasticity. Modifications in RNA repertoires are highly specific to the cues received by cells. In development, regional signals and transcription factors direct transcriptomic programs that specify cell types. However, even in post-mitotic cells, transcriptomes remain dynamic to drive structural and functional changes for plasticity. In particular, mature neurons – that integrate numerous and diverse cues - have developed cue-specific pathways for transcriptome remodeling to support various forms of plasticity (Greer and Greenberg, 2008). For example neuronal activity or growth factor signaling each trigger specific programs of de novo transcription resulting in the up-regulation of highly selective and specific sets of genes that modify neuronal wiring and function (Lambert et al., 2013; Mardinly et al., 2016; Russek et al., 2019; Spiegel et al., 2014).

More recent studies revealed that subcellular compartmentalization, in particular nuclear retention of mRNAs, represents another major mechanism to control functional gene expression. In non-neuronal cells, nuclear compartmentalization plays a substantial role in transcription noise buffering which prevents stochastic mRNA fluctuations in the cytosol (Bahar Halpern et al., 2015; Battich et al., 2015). Furthermore, active nuclear retention of mRNAs is emerging as a novel form of post-transcriptional gene regulation. Notably, nuclear retention of RNAs can be regulated through long-lasting processes such as neuronal differentiation (Yeom et al., 2021). Also, candidate gene approaches in several systems revealed that some stored and nuclear transcripts can be released into the cytosol upon acute signals, thereby rapidly increasing mRNAs’ availability for translation (Boutz et al., 2015; Mauger et al., 2016; Naro et al., 2017; Ninomiya et al., 2011; Prasanth et al., 2005). While further work is required to know whether this affects only rare RNAs or if this is a widespread mechanism, this discovery has generated considerable attention because it enhances functional gene expression independently of de novo transcription, a time-limiting step due to the finite processivity of the RNA polymerase II (Darzacq et al., 2007; Fuchs et al., 2014; Singh and Padgett, 2009; Tennyson et al., 1995; Veloso et al., 2014). The examination of transcript compartmentalization control is only emerging and further investigations are required to decipher its comprehensive potential in neuronal transcriptome remodeling. Notably, it remains unexplored whether this regulated cellular compartmentalization can also selectively remodel the transcriptome upon distinct signals.

Regulated intron retention (IR) has recently emerged as one candidate mechanism for nuclear retention and signaling-induced release of mRNAs. IR is a unique form of alternative splicing and consists of the persistence of a complete intron in otherwise fully synthetized mRNAs. IRs are highly prevalent and tightly regulated during development and in response to environmental signals supporting their major role in gene expression control (Jacob and Smith, 2017). IRs are a heterogeneous class of alternative splicing events that can direct their host mRNAs to multiple fates. A minority of intron retaining transcripts (IR-transcripts) are protein coding and exported to the cytoplasm where they generate protein isoforms (Grabski et al., 2021; Marquez et al., 2015). In other cases, IR elicits the degradation of the transcript either in the nucleus or the cytoplasm (Braunschweig et al., 2014; Wong et al., 2013; Yap et al., 2012). More recent studies shed light on IR as a mechanism for regulating transcriptome dynamics (Boutz et al., 2015; Gill et al., 2017; Mauger et al., 2016; Naro et al., 2017; Ninomiya et al., 2011; Park et al., 2017; Pendleton et al., 2017, 2018). Some IR-transcripts are initially targeted for nuclear retention where they remain stored in the nucleus for many hours and several days (Mauger et al., 2016; Naro et al., 2017). These transcripts form a reservoir of RNAs that can be released into the cytosol upon signals through splicing completion independently of new transcription. Interestingly, intron retention programs appear to target different sets of transcripts in different cellular systems. In neurons, an elevation of network activity and calcium influx has been shown to rapidly lead to the splicing completion of transcripts encoding proteins involved in cytoskeletal regulation and signaling pathways (Mauger et al., 2016). By contrast, in male gametes, some mRNAs coding for proteins implicated in spermatogenesis are subject to splicing completion in the latest stage of gametogenesis (Naro et al., 2017). The difference in the identity of regulated transcripts likely reflect cell class-specific IR programs. However, it remains unexplored whether stimuli trigger splicing completion of IR-transcripts by releasing a common brake of intron excision or whether there are cue-specific IR programs controlled through dedicated signaling pathways. Noteworthy, in yeast, different cellular stresses modify the splicing kinetics of distinct sets of constitutively spliced introns (Bergkessel et al., 2011). Moreover multiple signaling pathways have been implicated in regulation of alternative exon choices in response to external stimuli (Matter et al., 2002; Shin and Manley, 2004; Zhou et al., 2012). This raises the possibility that distinct cues may target select IR programs in other systems including mature neurons.

In the present study we systemically mapped the subcellular localization of neuronal IR-transcripts and their response to neuronal stimuli. We found that the majority of transcripts that stably retain introns are subject to nuclear retention. After neuronal stimulation, the vast majority of transcripts that complete splicing are exported to the cytosol indicating that IR is a widespread mechanism to control storage and on-demand release of mRNAs from the nucleus. Remarkably, stimulation with brain-derived neurotrophic factor *versus* a brief elevation of neuronal network activity mobilizes distinct pools of IR-transcripts. This cue-specificity of IR programs arises from the engagement of distinct signaling pathways that convey specific messages to the neuronal nucleus. Overall, we conclude that IR programs allow a rapid, transcription-independent and cue-specific remodeling of neuronal transcriptome during plasticity.

## Results

### The majority of stable intron-retaining transcripts are localized in the nucleus

To systematically assess sub-cellular localization of transcripts with stable intron retentions (IR), we developed a dedicated experimental workflow. We performed biochemical cell fractionation (Suzuki et al., 2010) and separated nuclear and cytosolic RNAs of mature mouse primary neocortical cells **(Figure 1A)**. To solely analyze the population of stable intron retaining transcripts (IR-transcripts) rather than transcripts containing transient IRs, we pharmacologically blocked transcription for 3 hours before collecting cells **(figure supplement 1A)**. For each sample: whole-cell extract, nuclear-enriched (designated as “Nucleus”) and cytosolic-enriched (designated as “Cytosol”) compartments, polyadenylated (poly(A)+) RNAs were isolated from three biological replicates and spike-in RNAs were added to assess the absolute nuclear-to-cytosol ratio of expressed transcripts **(see Materials and Methods)**. Samples were sequenced at high depth (ca. 100 million reads per sample, 100-mer reads). Ribosomal RNAs represented ca. 1% of the mapped reads **(Table 1)**, indicating that the enrichment of poly(A)+ RNAs was highly efficient.

**Figure 1.**
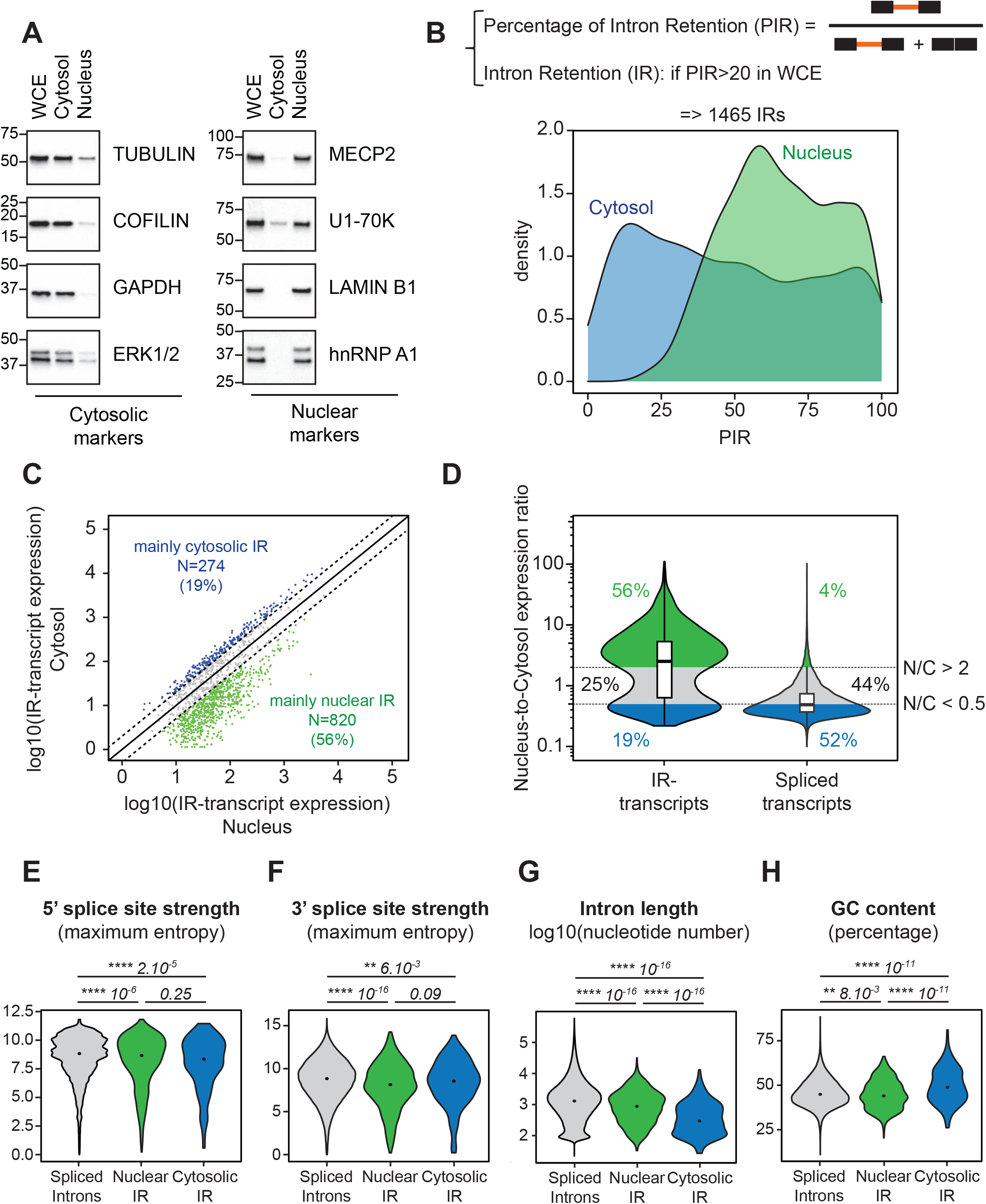
The majority of stable intron-retaining transcripts are localized in the nucleus. **(A)** Quality of the cell fractionation assays was controlled by western blot by assessing the distribution of nuclear and cytosolic markers in the whole cell extract (WCE), the cytosol and the nucleus. Protein lysates were isolated from mouse primary neocortical cells (14 days in culture) treated for 3hrs with the transcription inhibitor triptolide (1µM). (four independent cultures) **(B)** Top: percentage of intron retention (PIR) is assessed as the ratio between the IR-transcript expression and the total transcript expression (sum of intron-retaining and spliced transcripts). Intron are considered retained if minPIR ≥ 20 in WCE **(see Materials and Methods)**; 1465 IRs were then identified. Bottom: Density plot displaying the PIR distribution in cytosol (blue) and nucleus (green) of the 1465 retained introns. (three independent cultures) **(C)** Pairwise comparison of the expression of the 1465 intron-retaining isoforms (IR) in nuclear *versus* cytosolic fractions. IR-transcripts enriched in the nucleus are labeled in green (nuclear-to-cytosol expression ratio ≥ 2); IR-transcripts enriched in the cytosol are labeled in blue (nuclear-to-cytosol expression ratio ≤ 0.5). **(D)** Violin plots displaying the nucleus-to-cytosol expression ratio of the 1465 intron-retaining isoforms (left) and all expressed spliced transcripts (right). Percentage of nuclear-enriched (green) and cytosolic-enriched (blue) transcripts are indicated on the panel. **(E)** to **(H)** Violin plots displaying 5’ splice site strength (E), 3’ splice site strength (F), intron length (G) and GC content (H) of canonically spliced introns (grey), nuclear IRs (green) and cytosolic IRs (blue). The p-values calculated with a two-sided Mann-Whitney test are indicated on the top of each panel.

For every intron of the mouse genome, we analyzed the percentage of intron retention (PIR) with a previously established and validated pipeline (Mauger et al., 2016) **(see Materials and Methods, Figure 1B – figure supplement 1B)**. Introns were considered as retained if the PIR value was higher than 20 in whole-cell-extract. Similar to a previous analysis (Mauger et al., 2016), we identified 1465 stable IRs arising from 903 genes - among 10894 genes expressed in primary neocortical neurons. We then probed the distribution of intron retention levels for these events in each subcellular compartment **(Figure 1B)**. We found that PIR values were overall higher in the “Nucleus” than in the “Cytosol” (mean PIR_Nucleus_= 66; mean PIR_Cytosol_=48) suggesting that stable IR-transcripts are predominantly localized to the nucleus. To confirm this, we compared the expression of intron-retaining isoforms in the “Nucleus” and “Cytosol” samples **(Figure 1C and D)**. To calculate an absolute nuclear-to-cytosol ratio, we used the spike-in RNAs for normalizing the nuclear and cytosolic reads **(see Materials and Methods)**. Among the 1465 IR-transcript isoforms, 820 (56%) were strongly enriched in the “Nucleus” (nucleus-to-cytosol ratio>2), while 274 (19%) were more abundant in the “Cytosol” (nucleus-to-cytosol ratio<0.5); the remaining isoforms (25%) were similarly detected in the “Nuclear” and the “Cytosolic” fractions. By contrast, only a minority of fully spliced transcripts is enriched in the “Nucleus” (4%) and the majority of them is highly enriched in the “Cytoplasm” (52%). This indicates that the prevalence of IR-transcripts in the “Nucleus” samples is not a consequence of an inefficient biochemical fractionation **(Figure 1D)**.

To conclude, our data reveal an unprecedented large population of stable IR-transcripts predominantly localized in the nucleus and thus highlight that stable IR-transcripts are largely targeted for nuclear retention.

### Stable nuclear IRs share features with canonical spliced introns

We hypothesize that the nuclear localization of stable IR-transcripts is intronically encoded. To test this hypothesis, we examined whether the nuclear IRs harbor specific sequence properties. Nuclear retained introns exhibit weak 5’ and 3’ splice sites compared to canonically spliced introns - a general property of IRs (Boutz et al., 2015; Braunschweig et al., 2014; Mauger et al., 2016; Ullrich and Guigó, 2020; Yeom et al., 2021). However, the splice site strength of stable nuclear retained introns is indistinguishable from the one of cytosolic retained introns **(Figure 1E and F)**. Thus, the splice site sequences themselves are not dictating the nuclear localization of IR-transcripts.

However, stable nuclear retained introns display specific features in terms of length and GC content: while nuclear retained introns remain shorter than spliced introns, they are markedly longer than cytosolic retained introns **(Figure 1G)**. Similarly, GC content of nuclear retained introns is lower than the one of cytosolic retained introns and comparable to GC content of canonical spliced introns **(Figure 1H)**.

To conclude, in respect to several sequence features, stable nuclear IRs resemble canonically spliced introns and retention can be regulated by trans-acting factors. Thus, we hypothesize that a substantial proportion of stable nuclear retained introns preserves the ability to be excised through splicing; but as opposed to canonical spliced introns, enhancing cues may be required to promote their removal.

### Nuclear intron-retaining transcripts are regulated by several forms of neuronal stimulation

Intron retention rates have been reported to be regulated over days of neuronal differentiation (Braunschweig et al., 2014; Yap et al., 2012; Yeom et al., 2021) or in mature neurons in response to elevation of neuronal network activity (Mauger et al., 2016). Mature neurons exhibit specific forms of plasticity in response to specific plasticity cues. To explore whether such cues can acutely target subsets of IRs for rapid transcriptome remodeling, we first probed whether different stimuli can regulate IRs in mature neocortical neurons. We used two modes of neuronal stimulation. Mouse primary neocortical cultures were treated for one hour with i) bicuculline, an antagonist of GABA_A_ receptors which induces a robust increase in neuronal network activity, or with ii) the brain-derived neurotropic factor (BDNF) which is specifically released during forms of synaptic plasticity and facilitates long-term potentiation (Gottmann et al., 2009; Harward et al., 2016). In each condition, cells were treated with a transcription inhibitor to solely focus on IR-transcripts that are stable in unstimulated neocortical cells **(figure supplement 2A)**. Given that this study exclusively focuses on such stable IR-transcripts, we will simply designate them as “IR-transcripts” in the remainder of the manuscript. Both, bicuculline and BDNF stimulation induced a robust increase of ERK phosphorylation in nearly all neurons, indicating that both treatments stimulated the vast majority of neurons in culture **(Figure 2A and B – figure supplement 2B and C)**. Interestingly, we found that both bicuculline and BDNF stimulation regulate a sizeable set of IR-transcripts in the absence of de-novo transcription. Upon bicuculline treatment, the expression level of 430 IR-transcripts was altered (fold-change>20% and |z-score|>1.5); 382 IR-transcripts exhibit a lower expression upon stimulation (resulting from induced splicing or degradation, see Figure 3) and 48 transcripts were up-regulated (resulting from inhibition of basal splicing or enhanced stabilization, see Figure 3) **(Figure 2C – figure supplement 2D)**. As for BDNF stimulation, it regulated the expression of 385 IR-transcripts (243 down-regulated and 142 up-regulated) **(Figure 2D – figure supplement 2D)**. Importantly, on average regulated IR-transcripts are expressed at a similar level as unregulated IR-transcripts. Thus, these mRNAs constitute a major transcript pool rather than a lowly expressed subpopulation **(figure supplement 2E)**. The vast majority of regulated IR-transcripts were localized in the nucleus (nucleus-to-cytosol ratio>2; 84% and 79% of bicuculline- and BDNF-sensitive IR-transcripts respectively) **(Figure 2E and F)**. Remarkably, the population of regulated transcripts is even more enriched in the nucleus than the overall population of IR-transcripts (two-sided Mann-Whitney test, p-value<10^−5^ for both bicuculline- and BDNF-regulated transcripts) **(figure supplement 2F)**.

**Figure 2.**
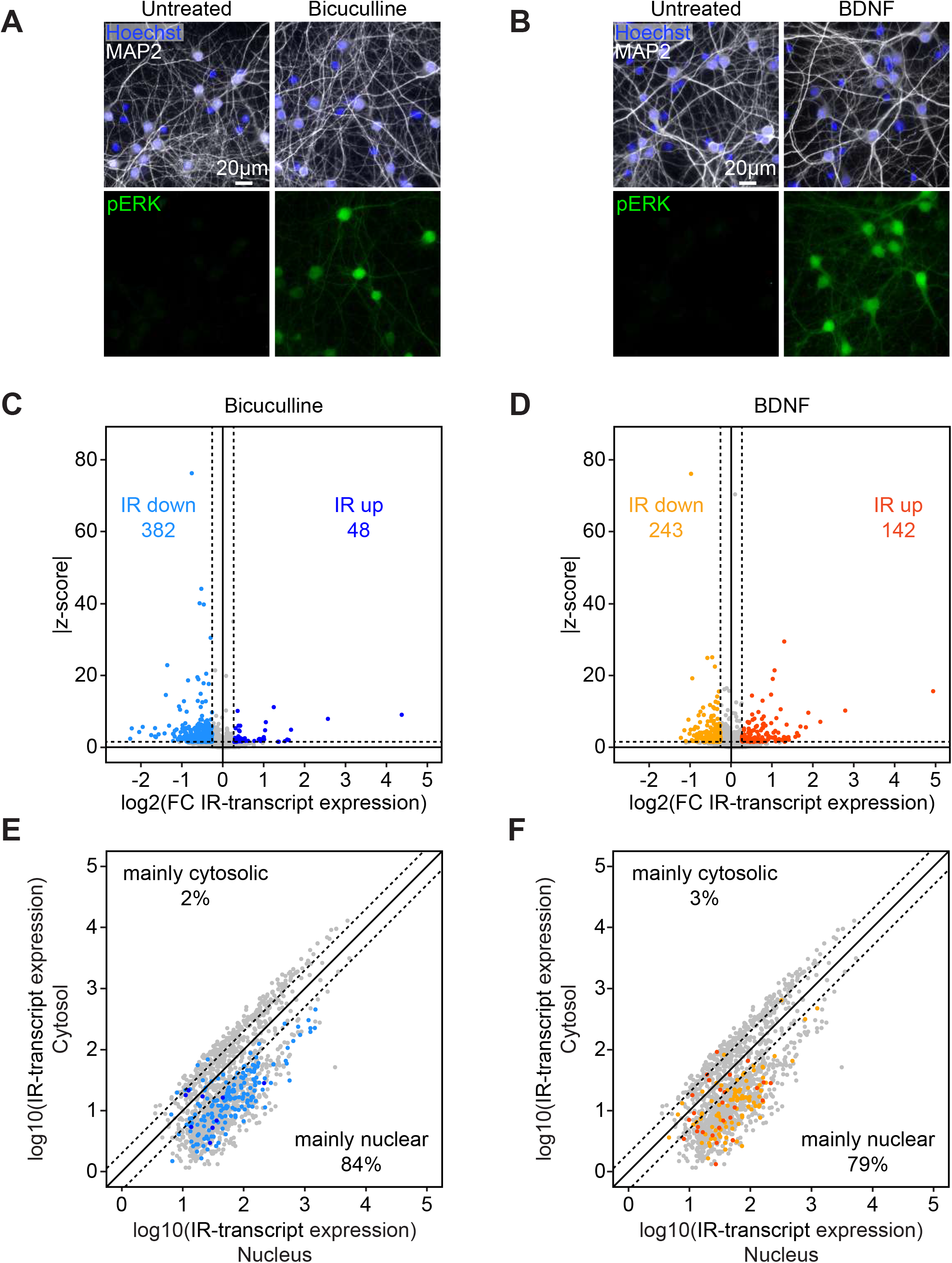
Nuclear intron-retaining transcripts are regulated by several forms of neuronal stimulation. **(A)** and **(B)** Efficiencies of bicuculline (A) and BDNF (B) stimulations were assessed by controlling the induction of ERK phosphorylation by immunostaining. Mouse primary neocortical cells (14 days in culture) were stimulated with bicuculline (20µM) or BDNF (50ng/mL) for 5min. Cells were stained with anti-phospho-ERK (green, bottom) and anti-MAP2 (white, top) antibodies and counterstained with Hoechst (blue, top). (four independent cultures) **(C)** and **(D)** Volcano plots displaying the expression fold change of IR-transcripts and the corresponding z-score (absolute value) upon bicuculline (C) and BDNF (D) stimulations. DRB (50µM) was applied to mouse primary neocortical cells for 2hrs; 1hr before cell collection, bicuculline (20µM) or BDNF (50ng/mL) were applied or not (control). Every transcript retaining an intron (minPIR ≥ 20%) in unstimulated condition were plotted. IR-transcripts were considered downregulated (light blue or orange) or upregulated (dark blue or red) if the following applied: fold change of IR-transcript expression ≥ 20% and |z-score| ≥ 1.5. (three independent cultures) **(E)** and **(F)** Pairwise comparison of the expression of regulated IR-transcripts in the nuclear and the cytosolic fractions of unstimulated mouse primary neocortical cells. Every transcript retaining an intron (minPIR ≥ 20%) in whole cell extract were plotted. Intron retaining transcripts down- or up-regulated upon bicuculline stimulation (E) are labeled in light blue and dark blue respectively and those down- or up-regulated upon BDNF stimulation (F) are labelled in orange and red respectively. (three independent cultures).

**Figure 3.**
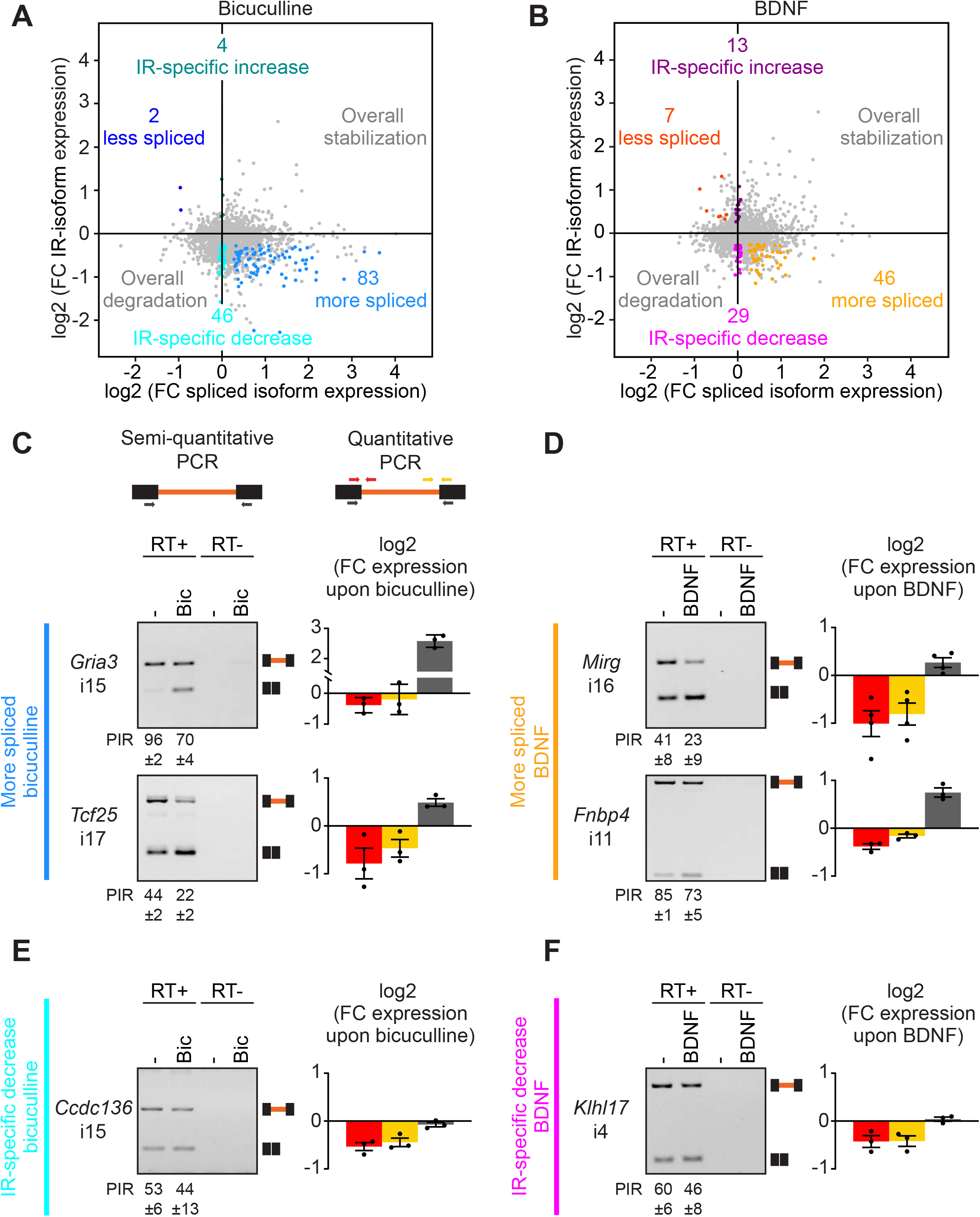
Neuronal stimulation regulates intron-retaining transcripts through splicing and degradation processes. **(A)** and **(B)** Pairwise comparison of the expression fold change (FC) of intron-retaining isoforms (IR) and the spliced isoforms upon bicuculline (A) or BDNF (B) stimulation. In all conditions, DRB (50µM) was applied to mouse primary neocortical cells (14 days in culture) for 2hrs; 1hr before cell collection, bicuculline (20µM) or BDNF (50ng/mL) were applied or not (control). Every transcript containing a retained intron (minPIR>20%) in control condition are plotted. IR-transcripts are considered regulated through splicing if the following applied: (1) IR-isoform expression fold change ≥ 20% and |z-score| ≥ 1.5, (2) spliced isoform expression fold change ≥ 20% and |z-score| ≥ 1.5, (3) expression of the IR-isoforms and the spliced isoforms evolved in opposite directions. IR-transcripts are considered regulated through degradation (IR-transcript specific decrease) if the following applied: (1) IR expression fold change ≥ 20% and |z-score| ≥ 1.5, (2) spliced expression fold change ≤ 5% and |z-score| ≤ 1. (three independent cultures) **(C)** to **(F)** RT-PCR validations of regulated IR-transcripts through splicing upon bicuculline stimulation (C) and BDNF stimulation (D) and through degradation upon bicuculline stimulation (E) and BDNF stimulation (F). Expression of the IR-isoforms and the spliced isoforms were analyzed by semi-quantitative PCR (left panels). Means and SEMs of PIR values are shown beneath each panel. In addition, fold changes in the expression of the IR-transcripts (red and orange) and spliced (dark grey) isoforms were assessed with real-time qPCR using three different primer sets, as represented in the top scheme. SEMs are displayed (three-four independent cultures). Note that the Gria3 spliced transcript (C) does not correspond to the canonical mRNAs and presumably arise from a first step of recursive splicing and thereby likely require splicing completion to generate fully mature Gria3 transcripts (Sibley et al., 2015).

Thus, our data reveals that multiple plasticity cues target nuclear IR-transcripts. We hypothesize that a large population of IRs can be regulated through nuclear processes including splicing.

### Neuronal stimulation regulates intron-retaining transcripts through splicing and degradation processes

The fate of IR-transcripts and their contribution to protein production is determined by the rates of splicing, degradation, and nuclear export. Each of these processes could be targeted for establishing specific transcriptome modifications in response to distinct forms of neuronal stimulation. Hence, before thoroughly probing the cue-specificity of IR programs, we dissected the contribution of degradation and splicing for the two forms of neuronal stimulation. We performed a pairwise comparison of the expression regulation of IR-transcripts and their counterpart spliced transcripts. Upon splicing, the decrease of intron-retaining isoforms is accompanied by an increase of the spliced isoforms. Applying stringent criteria to select IRs that follow this scheme **(see Materials and Methods)**, we found that 83 and 46 IR-transcripts underwent splicing upon neuronal stimulation with bicuculline and BDNF, respectively **(Figure 3A and B – figure supplement 3A and B)**. We performed targeted validations using semi-quantitative PCR and real-time quantitative PCR assays for several IR-transcripts. Notably, the transcripts encoding the AMPA receptor subunit GRIA3 and the transcription factor TCF25 exhibit a concomitant decrease of the intron-retaining isoforms and an increase of the spliced isoforms confirming their regulation through splicing induction **(Figure 3C – figure supplement 3C)**. We further validated a decrease of the intron-retaining isoforms and concomitant increase of the spliced isoforms of transcripts encoding the cytoskeletal regulator FNBP4 and the transcript arising from the microRNA containing gene Mirg **(Figure 3D – figure supplement 3D)**. Interestingly, we also identified IR-transcripts that showed increased retention and reduced levels of the spliced isoforms upon neuronal stimulation (2 for bicuculline, 7 for BDNF). This indicates that intron excision can also be slowed-down in response to signaling. Note that the apparent low number of IR-transcripts that undergo reduced splicing results from the fact we focused our analysis on IR-transcripts that were stable before stimulation; i.e., those associated with IRs whose retention level remains higher than 20% after transcription inhibition.

Remarkably, our data also reveal that a substantial population of regulated IRs cannot readily be explained by a splicing mechanism. Many IRs were associated with spliced and intron-retaining isoforms regulated in the same direction. They were either both increased or decreased suggesting a respective overall stabilization or degradation not instructed by IRs **(Figure 3A and B)**. We also found a sizeable set of regulated IR-transcripts whose spliced counterpart was not regulated **(see Materials and Methods)** suggesting that induced-degradation/stabilization process was specifically targeting the IR-transcript isoforms **(Figure 3A and B – figure supplement 3A and B)**. Note that in some cases, these events could also arise from a splicing reaction targeting other splice sites; however, our pipelines did not detect examples for such cases. Notably, 46 and 29 transcripts were destabilized upon stimulation with bicuculline and BDNF respectively. Interestingly, while our analysis focused on IR-transcripts that were stable in unstimulated conditions, we also found that a few (4 and 13) IR-transcripts were even more stable upon neuronal stimulation with bicuculline or BDNF. PCR assays confirmed the reliable identification of such regulation by degradation. For example, the transcript encoding the DNA double strand regulator CCDC136 exhibits a consistent decrease of intron-retaining isoforms but no change in the spliced isoforms in response to bicuculline stimulation **(Figure 3E – figure supplement 3C)**. Similar regulation was observed for the transcripts encoding the brain-specific actin regulator KLHL17 upon BDNF application **(Figure 3F – figure supplement 3D)**.

In aggregate, our data support that in mature neocortical neurons, neuronal signaling not only drives transcription-independent modifications of the neuronal transcriptome through splicing completion but also through transcript-specific degradation.

### Activity-dependent splicing of intron-retaining transcripts promotes cytosolic export of fully spliced transcripts

For a small number of selected IR-transcripts the neuronal activity-dependent splicing completion was shown to be followed by nuclear export and translation (Mauger et al., 2016). However, it remains unknown whether the release of nuclear retention of mRNAs upon splicing completion is a general mechanism. To address this question at a transcriptome-wide scale, we systematically probed the localization of IR-transcripts and their spliced mRNA counterparts in the nucleus and the cytosol 1-hour after bicuculline-mediated elevation of neuronal network activity. First, we mapped the total cellular repertoire of IRs and regulated IR-transcripts (whole cell extract samples, same criteria than in previous figures, **see Materials and Methods**). Nearly all regulated IR-transcripts were predominantly localized to the nucleus **(Figure supplement 4A)**. As expected, IR-transcripts regulated through splicing (based on whole cell extract data) were significantly less abundant in the nucleus in response to bicuculline stimulation (consistent with splicing being a nuclear process) **(Figure 4, left panel)**. A concomitant increase of the spliced isoforms was also observed in the nucleus while it was not the case for transcripts degraded upon stimulation **(Figure 4, middle panel)**. Remarkably, we found that the spliced transcripts were also significantly enriched in the cytosol one hour after bicuculline application indicating the newly spliced transcripts were exported to the cytosol after splicing completion **(Figure 4, right panel)**. By contrast, the spliced transcripts associated with IR-transcripts regulated through degradation were almost unchanged upon stimulation.

**Figure 4.**
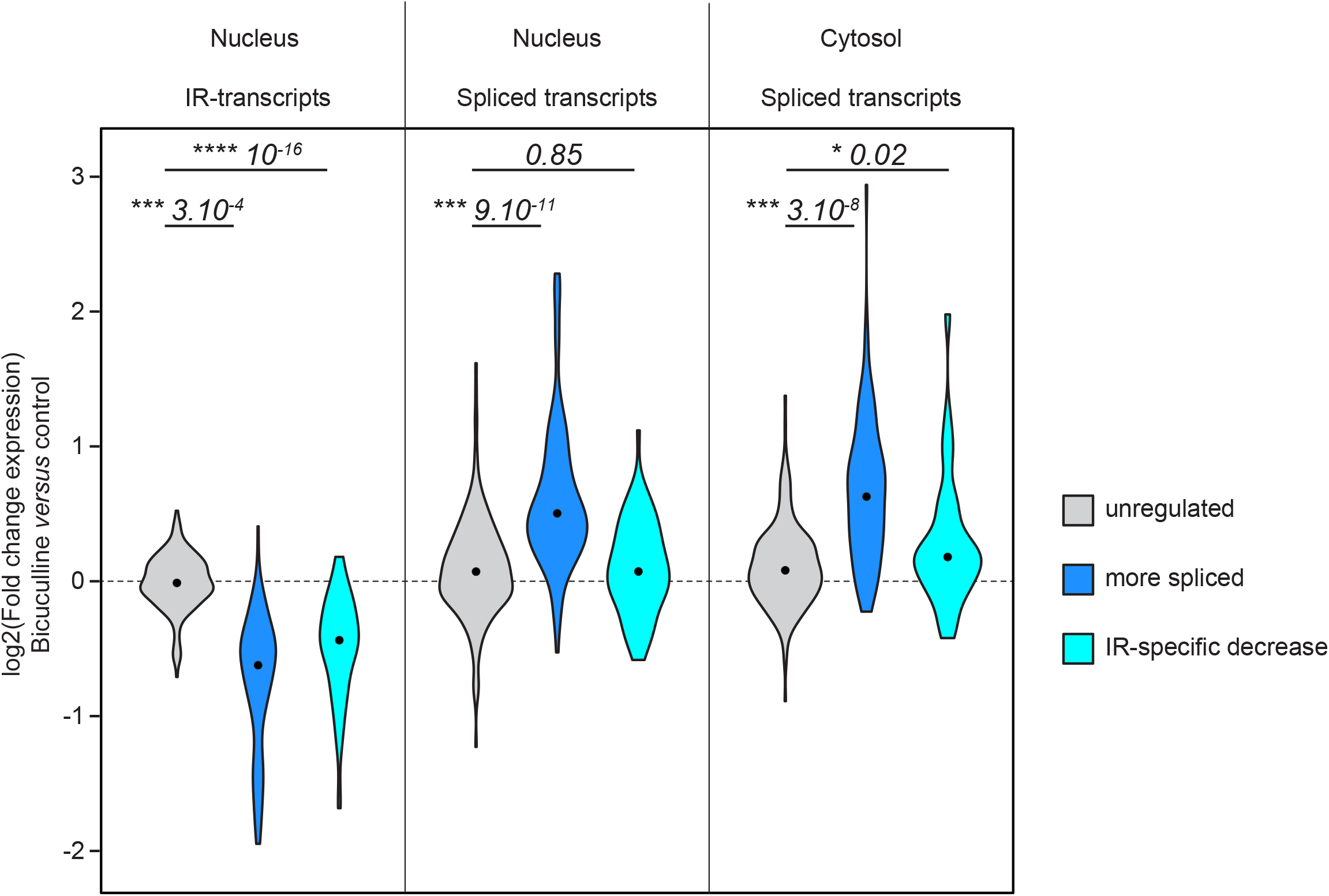
Activity-dependent splicing of intron-retaining transcripts promotes cytosolic export of fully spliced transcripts. Violin plots displaying expression fold change of IR-transcripts in the nucleus (left), spliced transcripts in the nucleus (middle) and spliced transcripts in the cytosol (right) upon bicuculline stimulation. Triptolide (10µM) was applied to mouse primary neocortical cells (14 days *in vitro*) for 3hrs; 1 hour before cell collection, bicuculline (20µM) was applied or not (control). Only transcripts that retained an intron in unstimulated and WCE conditions (minPIR ≥ 20%) were considered for the analysis. Transcripts were considered unregulated (grey) if the following applied: (1) fold change in the expression of the intron retaining transcripts ≤ 5% and |z-score| ≤ 1 in the WCE. Transcripts were considered regulated through splicing (dark blue) if the following applied: (1) IR-transcript expression fold change ≥ 20% and |z-score| ≥ 1.5 in the WCE, (2) Spliced isoform expression fold change ≥ 20% and |z-score| ≥ 1.5 in the WCE, (3) expression of the IR- and the spliced isoforms evolved in opposite directions in the WCE. Transcripts were considered regulated through degradation (IR-specific decrease, light blue) if the following applied: (1) IR-transcript expression fold change ≥ 20% and |z-score| ≥ 1.5 in WCE, (2) spliced expression fold change ≤ 5% and |z-score| ≤ 1 in WCE. The p-values calculated with a two-sided Mann-Whitney test are indicated on the top of the panel. (three independent cultures)

This suggests, that IR enables the nuclear compartmentalization of transcripts; furthermore, intron removal through splicing completion represents a widely-used mechanism for stimulus-dependent release of transcripts into the cytosol, thereby rapidly making them available for translation. Importantly, this major form of gene regulation occurs in the absence of alterations in total transcript levels **(Figure supplement 4B)** and, thus, is not detectable with conventional transcriptomic methods.

### Stimulus-specific regulation of sub-populations of intron retentions

Neurons undergo distinct forms of plasticity in response to specific cues. Thus, we then wondered whether the regulation of IR programs exhibits cue-specific mobilization of specific transcript pools. We compared the regulation of IR-transcripts upon stimulations with bicuculline and BDNF **(Figure 5A and B)**. This analysis clearly revealed three major categories of IR-transcripts: i) IR-transcripts that are regulated by both bicuculline and BDNF stimulation, ii) IR-transcripts that are solely regulated by bicuculline stimulation, and iii) IR-transcripts only affected by BDNF stimulation.

**Figure 5.**
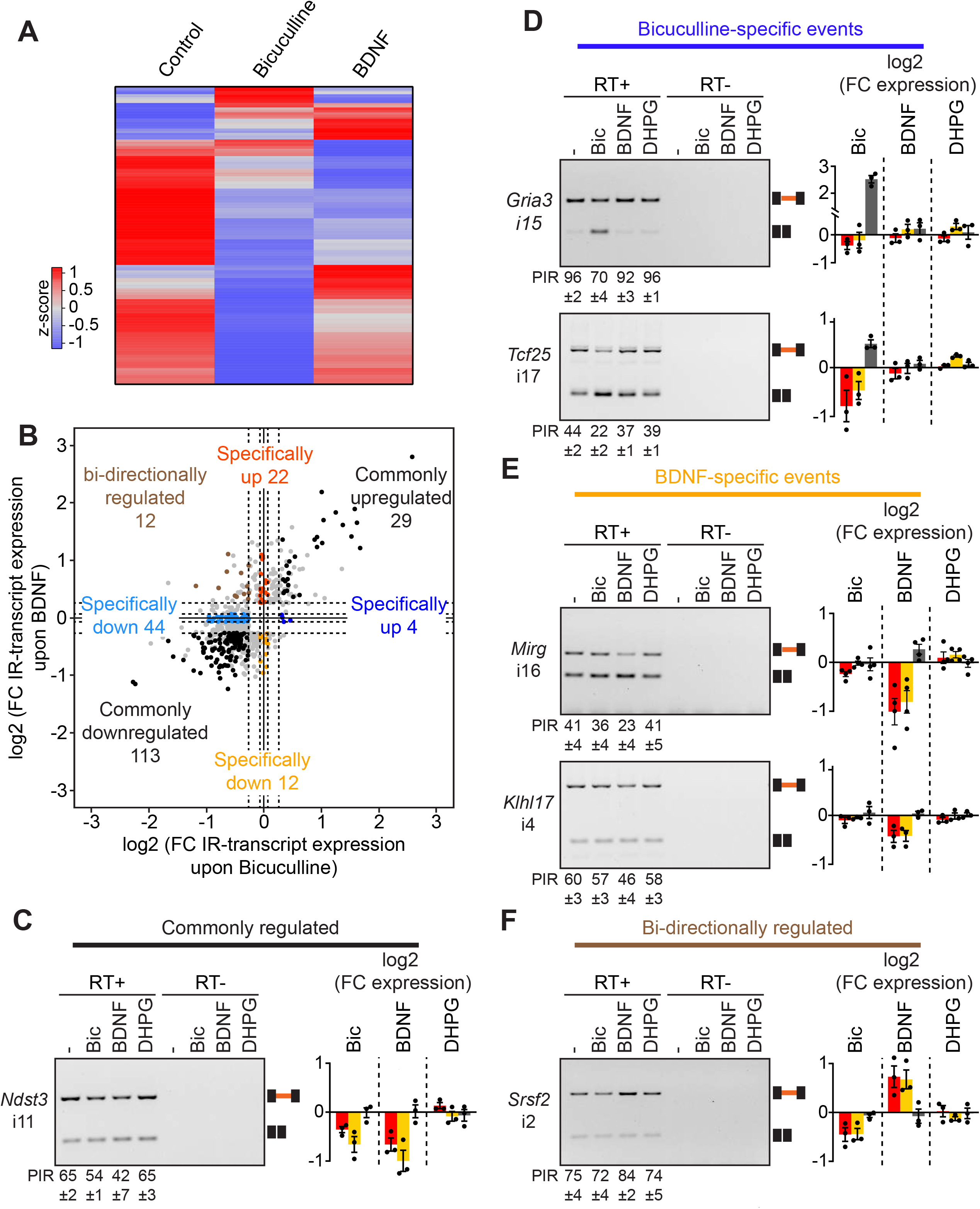
Stimulus-specific regulation of sub-population of intron retentions. **(A)** Heatmap of the IR-transcript expression values in control, bicuculline-stimulated and BDNF-stimulated conditions. IR-transcripts regulated in at least one condition are displayed (fold change of intron-retaining transcript expression ≥ 20% and |z-score| ≥ 1.5). **(B)** Pairwise comparison of the expression fold change (FC) of intron-retaining isoforms upon bicuculline and BDNF stimulation. Every transcript containing a retained intron (minPIR>20%) in control condition are plotted. IR-transcripts are considered regulated specifically upon bicuculline (light and dark blue) or (BDNF (orange and red)) stimulation if the following applied: (1) IR-transcript expression fold change ≥ 20% and |z-score| ≥ 1.5 upon bicuculline (or BDNF) stimulation (2) IR-transcript expression fold change ≤ 5% upon BDNF (or bicuculline) stimulation. IR-transcripts are considered commonly regulated upon bicuculline and BDNF stimulation (black) if the following applied: (1) IR-transcript expression fold change ≥ 20% and |z-score| ≥ 1.5 upon bicuculline stimulation (2) IR-transcript expression fold change ≥ 20% and |z-score| ≥ 1.5 upon BDNF stimulation (3) IR-transcripts evolved in the same direction upon bicuculline and BDNF stimulations. IR-transcripts are considered bi-directionally regulated upon bicuculline and BDNF stimulations (brown) if the following applied: (1) IR-transcript expression fold change ≥ 20% and |z-score| ≥ 1.5 upon bicuculline stimulation (2) IR-transcript expression fold change ≥ 20% and |z-score| ≥ 1.5 upon BDNF stimulation (3) IR-transcripts evolved in opposite directions upon bicuculline and BDNF stimulations. **(C)** to **(F)** RT-PCR validations of IR-transcripts commonly regulated (C), specifically regulated upon bicuculline stimulation (D), specifically regulated upon BDNF stimulation (E), and bi-directionally regulated (F). Expression of the IR-transcripts and the spliced isoforms were analyzed by semi-quantitative PCR (left panels). Means and SEMs of PIR values are shown beneath each panel. In addition, fold changes in the expression of the IR-transcripts (red and orange) and spliced isoforms (dark grey) were assessed with real-time qPCR. SEMs are displayed (three-four independent cultures).

For the category of commonly regulated IR-transcripts, 113 and 29 IR-transcripts were respectively down- and up regulated by both bicuculline and BDNF stimulations **(Figure 5B)**. For instance, the transcript encoding the metabolic enzyme NDST3 associated with schizophrenia is regulated upon both bicuculline and BDNF stimulation **(Figure 5C – figure supplement 5A and B)**. Nevertheless, *Ndst3* regulation harbors specificity as another stimulus (the group I metabotropic glutamate receptor agonist DHPG) did not impact its IR profile **(Figure 5C)**. Amongst commonly regulated transcripts, we could identify with high confidence 12 IR-transcripts regulated by splicing and 4 transcripts regulated by degradation upon both stimuli. Note that because we used very stringent criteria to identify transcripts regulated through splicing *versus* degradation, we could not confidently assign many of regulated IR-transcripts - identified in **Figure 3** - to splicing or degradation.

Remarkably, a sizeable population of IR-transcripts were specifically regulated by only one mode of stimulation **(Figure 5A and B)**. Using stringent criteria (fold-change < 5% for unregulated events, **see Materials and Methods**), we confidently identified 48 and 34 IR-transcripts solely regulated upon bicuculline or BDNF stimulation, respectively. Notably, the IR-transcripts encoding the AMPA receptor subunit GRIA3 and the transcription factor TCF25 are only regulated upon bicuculline stimulation but do not exhibit any change in response to upon BDNF application **(Figure 5D – figure supplement 5A and B)**. To further probe the selective regulation of these targets we used the group I mGluR agonist DHPG and similarly found no change in IR in these transcripts **(Figure 5D)**. By contrast, the transcripts arising from the miRNA-containing gene Mirg and the transcripts encoding the brain-specific actin regulator KLHL17 are exclusively regulated by BDNF stimulation but not upon stimulation with bicuculline or DHPG **(Figure 5E – figure supplement 5A and B)**.

Another striking category of specific IRs is associated with IR-transcripts that are bi-directionally regulated upon bicuculline *versus* BDNF stimulation. More precisely, some cue-specific IR-transcripts are regulated upon both bicuculline and BDNF stimulations but in opposite directions **(Figure 5E)**. For instance, the IR-transcripts encoding the splicing factor SRSF2 undergo degradation upon stimulation with bicuculline, while BDNF signal stabilized them **(Figure 5F – figure supplement 5B)**.

In sum, our data reveals that regulated IR-transcripts can be subdivided in several classes: some IRs are commonly regulated by bicuculline and BDNF stimulation whereas others are cue-specific. Thus, targeting IRs represents a way to specifically remodel the neuronal transcriptome upon distinct forms of neuronal stimulation.

### Stimulation-specificity of intron retention programs is conveyed by distinct signaling pathways

To obtain insight into the mechanism of neuronal cue-specific regulation of IR, we probed the signaling pathways involved in stimulated intron excision. Neuronal activity-dependent signaling elicited by elevation of network activity by bicuculline treatment relies on calcium signaling either through NMDA receptors or voltage-dependent calcium channels. As for BNDF-signaling events, they largely depend on the mitogen-activated protein kinase (MAPK) pathway. We thus wondered whether the stimulation-specificity of IR programs is conveyed by the differential activation of calcium signaling and MAPK pathways.

Remarkably, the application of the selective NMDA receptor antagonist AP5 impaired the bicuculline-dependent intron excision of *Tcf25* and *Gria3* transcripts **(Figure 6A – figure supplement 6A)**. Intron excision in these transcripts was also suppressed by pharmacological inhibition of calcium^2+^/calmodulin-dependent protein kinase (CaMK) - a downstream pathway activated by NMDAR-dependent calcium entry. By contrast, the MAPK antagonist U0126 did not impact splicing induction of bicuculline-sensitive transcripts. Conversely, for BDNF-sensitive introns, we found that the MAPK pathways is essential for the splicing of *Mirg* and the stabilization of *Srsf2* IR-transcripts upon BDNF stimulation **(Figure 6B – figure supplement 6B)**. However, the pharmacological inhibition of NMDA receptors and CaMK pathways did not preclude their regulation upon BDNF stimulation. Overall, these results uncover a signaling-pathway specificity of IR programs.

**Figure 6.**
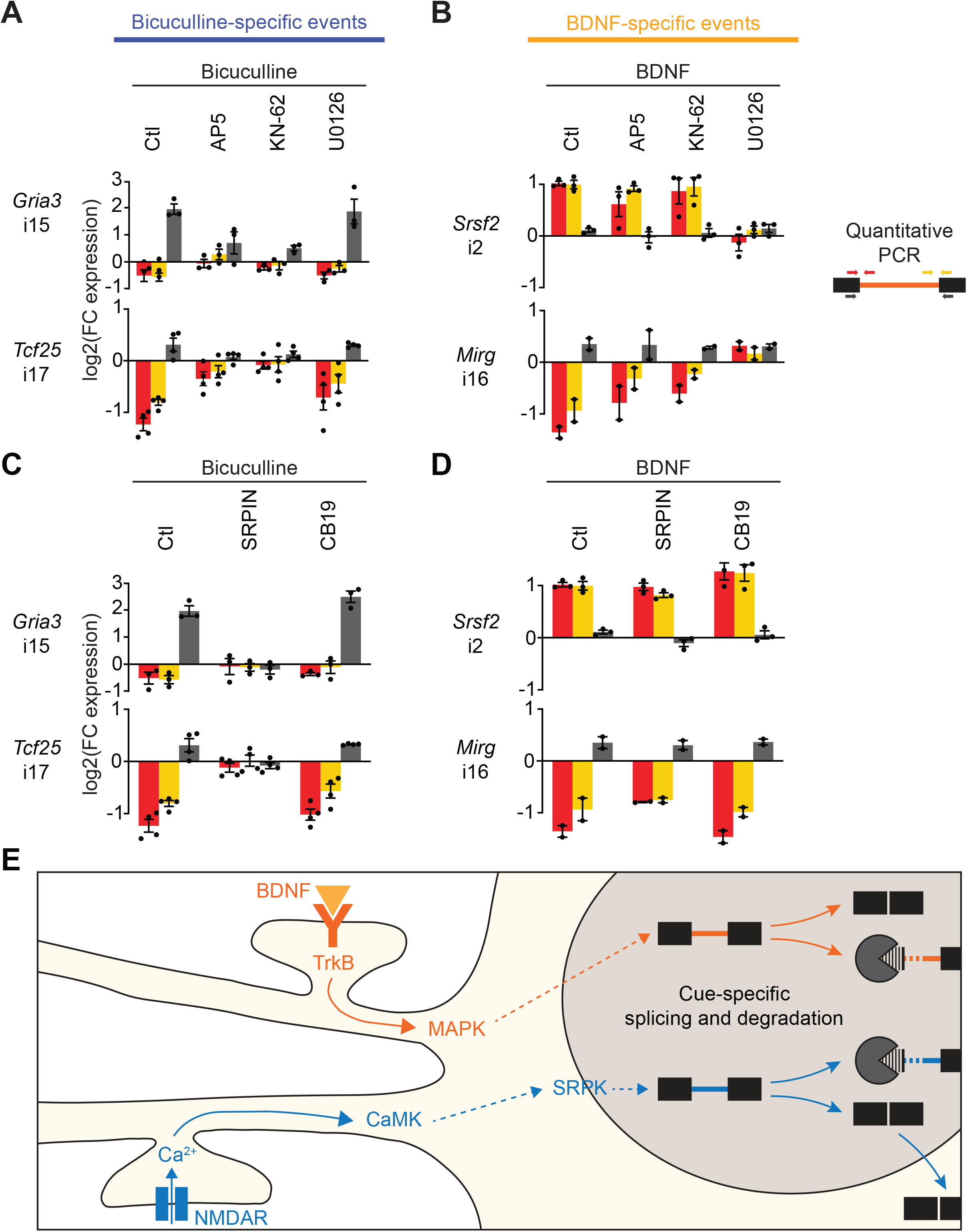
Stimulation-specificity of intron retention programs is conveyed by distinct signaling pathways. **(A)** and **(B)** Analysis of signaling pathways involvement in the regulation of IR-transcripts upon bicuculline (A) and BDNF (B) stimulations. In all conditions, DRB (50µM) was applied to mouse primary neocortical cells (14 days in culture) for 2hrs; 1hr before cell collection, bicuculline (Bic, 20µM) or BDNF (50ng/mL) was added to the cultures. 15min before bicuculline or BDNF stimulation, different pharmacological treatments were applied in order to block NMDA receptors with the antagonist AP5 (50µM) or the Calcium^2+^/calmodulin-dependent protein kinases (CaMK) with KN-62 (10µM) or the mitogen-activated protein kinases (MAPK) with U0126 (10µM). Fold changes in the expression of the IR-(red and orange) and spliced (dark grey) isoforms were assessed with real-time qPCR using three different primer sets, as represented in the right scheme. SEMs are displayed (two-to-four independent cultures). **(C)** and **(D)** Analysis of SPRK and CLK involvement in the regulation of IR-transcripts upon bicuculline (C) and BDNF (D) stimulation. In all conditions, DRB (50µM) was applied to mouse primary neocortical cells (14 days in culture) for 2hrs; 1hr before cell collection, bicuculline (Bic, 20µM) or BDNF (50ng/mL) was added to the cultures. 15min before bicuculline or BDNF stimulation, different pharmacological treatments were applied in order to block SR-protein kinases (SRPK) with SRPIN340 (10µM) or CDC2-like kinases (CLK) with KH-CB19 (10µM). Fold changes in the expression of the IR-(red and orange) and spliced (dark grey) isoforms were assessed with real-time qPCR using three different primer sets, as represented in the right scheme. SEMs are displayed (two-to-four independent cultures). **(E)** Working model. Selective IR programs – elicited upon distinct forms of neuronal stimulations and the subsequent activation of signaling pathways – remodel neuronal transcriptome in a cue-specific manner.

To obtain deeper insight into this cue-specific regulation, we focused on kinases previously implicated in signaling-dependent alternative splicing control (Shin and Manley, 2004). SR-protein kinases (SRPK) and the CDC2-like kinases (CLK) families have previously been shown to link external cues and alternative splicing regulation in non-neuronal cells (Ninomiya et al., 2011; Zhou et al., 2012). These kinase families both regulate the phosphorylation status - and consequently the activity - of SR proteins, the main family of splicing factors. Interestingly, SRPIN340, a pharmacological inhibitor of SRPK impeded intron excision of *Tcf25* and *Gria3* transcripts upon bicuculline treatment **(Figure 6C)**. However, SPRK inhibition had no effect on the BDNF-dependent splicing of *Mirg* or the BDNF-dependent regulation of *Srsf2* transcripts **(Figure 6D)**. Interestingly, KH-CB19, a pharmacological blocker of CLK kinase activity had no impact on neither bicuculline- or BDNF-dependent introns. Thus, these results further support that the cue-specificity of IR programs strongly relies on the activation of selective signaling pathways transduced to the nucleus to convey the mobilization of distinct sets of IR-transcripts.

In summary, this work identifies that selective IR programs elicited upon distinct forms of neuronal stimulation. The activation of distinct signaling pathways drives the remodeling of the neuronal transcriptome in a cue-specific manner **(Figure 6E)**.

## Discussion

In this work, we revealed that regulated intron retention is a widespread, cue-specific mechanism for neuronal transcriptome remodeling. We performed a comprehensive mapping of the subcellular localization of mRNAs in mature neurons and revealed that transcripts that stably retain introns are broadly targeted for nuclear retention. We systematically probed these transcripts upon neuronal stimulation and found that sub-populations of nuclear-retained transcripts are bi-directionally regulated in response to several cues: some appear targeted for degradation while others undergo splicing completion to generate fully mature mRNAs. This latter set of transcripts is exported to the cytosol to increase functional gene expression. Remarkably, distinct groups of IR-transcripts are regulated depending on the form of stimulation and this selectivity arises from the activation of specific signaling pathways. Overall, our data identifies reversible IR as a major regulator of nuclear mRNA retention and cue-specific mobilization

### Targeting intron retentions is a widespread mechanism to acutely regulate subcellular compartmentalization of transcripts upon cellular signals

In the present study, we show that retention of select introns in mRNAs is a widespread mechanism to control transcript localization. The nuclear localization of transcripts that stably retain select introns makes them unavailable for protein synthesis **(Figure 1)**. Previous work localized instable IR-transcripts to the nucleus (Braunschweig et al., 2014). Here, we systemically mapped the subcellular localization of stable IR-transcripts, and find this population to be abundant in the nucleus of mature neurons **(Figure 1)**. This is consistent with previous candidate gene studies in other cell classes, including fibroblastic cells, stem cells and male gametes, which reported nuclear localization of IR-transcripts. (Boutz et al., 2015; Mauger et al., 2016; Naro et al., 2017; Ninomiya et al., 2011).

We hypothesized that controlling intron retention and excision patterns represents a general mechanism to mobilize specific mRNA pools for functional gene expression. In line with this, we shed light on the extensive export of transcripts that undergo splicing completion in response to neuronal stimulation **(Figure 4)**. Other studies also revealed a link between IRs and the nucleo-cytosolic localization of their host transcripts (Boutz et al., 2015; Mauger et al., 2016; Naro et al., 2017; Ninomiya et al., 2011; Yeom et al., 2021). Though, in most of the cases, it was unclear i) whether existing transcripts repurposed their fate and localization through splicing completion or ii) whether co-transcriptional production of spliced isoforms is required for their cytosolic expression. Indeed, the fate and nuclear localization of some IR-transcripts can be irreversibly set up from their transcription (Pendleton et al., 2018). By contrast, we show here that a sizeable population of IR-transcripts can repurpose their fate upon environmental signals and promote their release into the cytosol. Such a transcription-independent mechanism likely evolved as it facilitates rapid remodeling of the transcriptome, independently of transcription which is time-limiting due to the finite processivity of the RNA-polymerase II (Darzacq et al., 2007; Fuchs et al., 2014; Singh and Padgett, 2009; Tennyson et al., 1995; Veloso et al., 2014)

### Intron retention programs remodel the neuronal transcriptome in a cue-specific manner

Previous studies showed that IR profiles across tissues and cell types are modified by numerous signals including cellular differentiation, neuronal stimulation, metabolic homeostasis and cellular stress (Boutz et al., 2015; Green et al., 2020; Haltenhof et al., 2020; Mauger et al., 2016; Naro et al., 2017; Park et al., 2017; Parra et al., 2018; Pendleton et al., 2018; Pimentel et al., 2016; Quesnel-Vallières et al., 2016; Wong et al., 2013). In the present work, we unveil that in mature neurons, IR programs are cue-specific and that selectivity arises from the activation of distinct neuronal signaling pathways **(Figure 5 and 6)**. Rather than constituting a universal program, distinct subsets of IR-transcripts are regulated in response to specific neuronal stimuli **(Figure 5)**. This specificity of IR programs in neocortical cells suggests that they contribute to exquisite neuronal plasticity events. Interestingly, in Drosophila, learning paradigms elevate the spliced isoform of *Orb2A* - encoding a protein involved in memory consolidation - over an unspliced isoform that is predominant in naive flies (Gill et al., 2017). The novel insights into the signaling mechanisms of neuronal IR regulation uncovered here will pave the way to probing the contribution of IR programs in transcription-independent plasticity events *in vivo*.

## Materials and Methods

### Primary neocortical cultures and pharmacological treatments

All procedures related to animal experimentation were reviewed and approved by the Kantonales Veterinäramt Basel-Stadt. Dissociated cultures of neocortical cells were prepared from E16.5 mouse embryos (embryonic stage 16.5). Neocortices were dissociated by the addition of papain (130 units, Worthington Biochemical, LK003176) for 30 min at 37°C. Cells (45,000 cells/cm2) were maintained in neurobasal medium (Gibco, 21103-049) containing 2% B27 supplement (Gibco, 17504-044), 1% Glutamax (Gibco, 35050-61), and 1% penicillin/streptomycin (Bioconcept, 4-01F00-H) for 14 days. The following reagents for pharmacological treatments were used (at the indicated concentrations and from the stated sources): DRB (50 µM, Sigma, D1916), triptolide (1 µM, Sigma, T3652), bicuculline (20 µM, Tocris, 0130), BDNF (50 ng/mL, Sigma, B7395), DHPG (10 µM, Tocris, 0805), DL-AP5 (50 µM, Tocris, 3693), KN-62 (10 µM, Tocris, 1277), U0126 (10µM, Tocris, 1144), SRPIN340 (10 µM, Tocris, 5063) and KH-CB19 (10 µM, Tocris, 4262).

### Cellular fractionation

Cell fractionation experiments were performed according the protocol from Suzuki et al. (Suzuki et al., 2010). Briefly, two million cells plated in a 10-cm^2^ dish were collected in ice-cold PBS. After 10 sec-centrifugation, the supernatant was removed from each sample and the cell pellet was resuspended in 450 μL ice-cold 0.1% NP40 in PBS. One aliquot was collected as the whole-cell extract and then the leftover was spun for 10 sec. The supernatant was collected as the cytosolic-enriched fraction and the pellet (after one wash with 450 μL 0.1% NP40 in PBS) as the nuclear-enriched fraction.

### Western blot and antibodies

Total proteins were separated by electrophoresis on 4%–20% gradient PAGE gels (Bio-Rad, 4561093) and transferred onto nitrocellulose membrane. The following antibodies were used: anti-phosphoCREB (Ser133) (Millipore, aa77-343), anti-CREB (clone 48H2, Cell signaling, 9197), anti-phosphoERK (Cell signaling, 4370S), anti-ERK (Cell signaling, 4695S), anti-phospho-CTD RNA polymerase II (Ser2) (Abcam, ab5095), anti-CTD RNA polymerase II (Abcam, ab817), anti-N-terminal RNA polymerase II A-10 (Santa Cruz, sc17798), anti-ACTININ (Abcam, ab68194), anti-COFILIN (Abcam, ab54532), anti-GAPDH (clone D16H11, Cell Signaling, 5174), anti-MECP2 (Cell Signaling, 3456), anti-U1-70K (clone H111, Synaptic Systems, 203011), anti-LAMIN B1 (Abcam, ab133741), anti-hnRNPA1 (clone D21H11, Cell Signaling, 8443), anti-betaIII TUBULIN (Abcam, ab18207).

### Immunocytochemistry and image analysis

For immunocytochemistry, mouse primary neocortical neurons were fixed with 4% PFA in 1X PBS for 15min. Cells were then permeabilized with ice-cold methanol for 10min and blocked (5% donkey serum, 0.03% Triton X-100 in 1X PBS) for 1hr at room temperature. Primary antibody incubation was performed overnight at 4°C in a humidified chamber. Secondary antibody incubation was then performed for 1hr at room temperature. The following antibodies were used: anti-phosphoERK (Cell signaling, 4370S), anti-MAP2 (Synaptic systems, 188004), Cy5-conjugated donkey anti-guinea pig (Jackson, 706-715-148) and Cy3-conjugated donkey anti-rabbit (Jackson, 711-165-152). Imaging was performed on a widefield microscope (FEI MORE) with a 40X objective (NA 0.95, air). Image analysis was performed on Fiji (Schindelin et al., 2012). Briefly, a mask for MAP2 signal was created and neuronal cell bodies were manually delimited. The mean of phospho-ERK signal intensity for each neuronal cell body was then measured.

### RNA isolation and reverse transcription

Cells were lysed using Trizol reagent (Sigma, T9424). Total RNAs were isolated and DNase treated on columns (RNeasy micro kit, Qiagen, 74004) following the manufacturer’s instructions. The cDNA libraries were built using between 100 and 200ng RNA reverse transcribed with SuperScript III reverse transcriptase (Thermo Fisher, 18080044), dNTPs (Sigma, D7295) and oligo(dT)_15_ primer (Promega, C1101).

### PCR

Semi-quantitative PCR was performed using Phusion High-Fidelity DNA Polymerase (New England Biolabs, M0530L) and revealed with GelRed (Biotium, 41003). For each PCR, the number of cycles necessary to end the amplification in its exponential phase was determined. In the case of long introns (>1,000–1,500 bp), multiplex PCRs were performed to amplify both the intron-retaining and the spliced transcript isoforms (i.e., using three primers: one primer complementary to an internal region of the intron in addition to the primers mapping to each flanking exon).

Real-time quantitative PCRs were performed with FastStart Universal SYBR GreenMaster (Roche, 04-913-850-001). PCRs were carried out in a StepOnePlus qPCR system (Applied Biosystems) with the following thermal profile: 10 min at 95°C, 40 cycles of 15s at 95°C and 1 min at 60°C. Real-time quantitative PCR assays were analyzed with the StepOne software. The primers used for PCRs are listed in **Table 2**.

### Deep RNA-sequencing

RNA samples were quality-checked on the TapeStation instrument (Agilent Technologies) using the RNA ScreenTape (Agilent, 5067-5576) and quantified by Fluorometry using the QuantiFluor RNA System (Promega, E3310). Library preparation was performed, starting from 200ng total RNAs, using the TruSeq Stranded mRNA Library Kit (Illumina, 20020595) and the TruSeq RNA UD Indexes (Illumina, 20022371). 15 cycles of PCR were performed. Libraries were quality-checked on the Fragment Analyzer (Advanced Analytical) using the Standard Sensitivity NGS Fragment Analysis Kit (Advanced Analytical, DNF-473) revealing an excellent quality of libraries (average concentration was 158±20 nmol/L and average library size was 351±6 base pairs). Samples were pooled to equal molarity. The pool was quantified by Fluorometry using the QuantiFluor ONE dsDNA System (Promega, E4871). Libraries were sequenced Paired-End 101 bases using the HiSeq 2500 or NovaSeq 6000 instrument (Illumina) and the S2 Flow-Cell loaded. Details on the number of sequenced reads for each sample are given in **Table 1**. Quality control of read sequences was performed in collaboration with the company GenoSplice technology (http://www.genosplice.com). Confidence score per base and per sequence, GC content, sequence length, adapter content and presence of overrepresented sequences was assessed using FAST-QC.

### RNA-sequencing read alignment

For the RNA-sequencing data analysis, reads were aligned onto the mouse genome assembly mm10 using the STAR aligner version 2.4.0f1 (Dobin et al., 2013). The following parameters were used: –outFilterMismatchNmax 2 and – outFilterMultimapNmax 1 to filter out reads that have more than two mismatches and reads that map to multiple loci in the genome. Moreover, reads with <8 nt overhang for the splice junction on both sides were filtered out using –outSJFilterOverhangMin 30 8 8 8. Then, read alignment files (bam) were processed with custom Perl scripts using the library Bio::DB::Sam. For each segment comprising a pair of consecutive exons (exon1 and exon2) and the intermediate intron, reads that mapped (1) exon1-intron junctions, (2) intron-exon2 junctions (3), exon1-exon2 junctions and (4) introns were counted. Note that the data discussed in this manuscript will be deposited in NCBI’s GeneExpression Omnibus (Edgar et al., 2002) before publication.

### PIR, intron-retaining isoform expression and spliced isoform expression

For each segment in the mouse genome, comprised of a pair of consecutive exons and the intervening intron annotated in FastDB (http://www.easana.com) (de la Grange et al., 2005), we assessed the expression level of transcripts retaining the intron (IR-isoforms) as the average number of reads mapping the 5’ and 3’ exon-intron junctions (= mean(cov(E1I), cov(IE2)). The expression level of the spliced isoform was estimated as the number of exon-exon junction reads (= cov(E1E2)). Given that coverage (cov) is by definition normalized by the length of the analyzed sequence, cov is corresponding to the absolute number of reads mapping a junction. All the read coverage values were normalized by the number of total mapped reads for each individual sample. Segments were considered as non-expressed and filtered out if (cov(E1E2) + mean(cov(E1I), cov(IE2)) < 10 in at least one sample. Note that, for the cell fractionation samples, the segment expression filter only applied to WCE condition. Also, segments whose spliced isoforms was not significantly expressed in any of the compared samples were filtered out: (cov(E1E2) < 10). Again, for cell fractionation samples, this filter only applied to WCE condition. The PIR was then calculated as the expression level of the IR-isoforms over the sum of the expression levels of the IR-isoforms and the spliced isoforms (PIR = mean(cov(E1I), cov(IE2))/(cov(E1E2) + mean(cov(E1I), cov(IE2))). We defined introns as retained if their PIR exceeded 20 and if they fulfilled the following criterion: the read coverage along the entire intron must represent at least 20% of the sum of the expression of the spliced and IR-isoforms (minPIR = (min(cov(E1I), cov(IE2), cov(I))/(cov(E1E2) + mean(cov(E1I), cov(IE2); min = minimum). In this way, we ensured that reads mapping the exon-intron junctions were indeed due to IR rather than other events, such as the usage of alternative 5’ or 3’ splice sites.

### Analysis of intron-retaining transcript subcellular localization and spike-in RNAs

Because the quantity of RNAs in the “Nucleus” and the “Cytosol” fractions are not equal, the real enrichment of a given transcript in the nucleus *versus* the cytosol is lost during the RNA sequencing library preparation. To overcome this hurdle, we added a defined amount of spike-in RNAs SIR-Set3 (Lexogen, Iso Mix E0/ERCC #051) in each fraction arising from a constant number cells before adjusting the RNA quantities during library preparation. Note that we put twice less spike-in RNAs for the nuclear samples. For our analysis, we only considered ERCC that are covered by >10 reads in every sample (18 ERCCs). We then calculated the cytosol-to-nucleus abundance ratio of each ERCCs. A correction factor was then assessed as the mean of cytosol-to-nucleus ratio for all ERCCs (1.03 ±0.07). Given that we introduce twice less spike-ins in the nuclear samples, the final correction factor used in this study is 0.515 (=1.03/2). We applied this factor to the RNA sequencing data to calculate a true nucleus-to-cytosol enrichment. IR-transcripts were considered enriched i) in the nucleus if (mean IR-isoform expression_nucleus_ / mean IR-isoform expression_cytosol_ ≥ 2) or ii) in the cytosol if (mean IR-isoform expression_nucleus_ / mean IR-isoform expression_cytosol_ ≤ 0.5). For the evaluation of splice site strength, maximum entropy scores for 9-bp 5’ splice sites and 20-bp 3’ splice sites were calculated using MaxEntScan (Yeo and Burge, 2004).

### Analysis of stimulation-dependent intron-retaining transcripts

Intron-retaining transcripts were considered as regulated if the following criteria applied: (1) the fold change mean of IR-transcript expression (FC IR expression) ≥ 20%, and (2) |z-score| of IR-transcript expression (|z-score|_IR expression_) ≥ 1.5, where z-score = (var(IR-transcript expression_unstimulated_) + var(IR-transcript expression_stimulated_)) – covar(IR-transcript expression)).

For all regulated intron-containing transcripts, the stimulation-specificity was defined as follow: <u>commonly regulated IRs</u>: (1) FC IR expression ≥ 20% upon both bicuculline and BDNF stimulations, (2) IRs regulated in the same direction upon both bicuculline and BDNF stimulations, (3) |z-score|_IR expression_ ≥ 1.5 upon both bicuculline and BDNF stimulations; <u>differentially regulated IRs</u>: (1) FC IR expression ≥ 20% upon both bicuculline and BDNF stimulations, (2) IRs regulated in opposite directions upon bicuculline and BDNF stimulations, (3) |z-score|_IR expression_ ≥ 1.5 upon both bicuculline and BDNF stimulations; <u>bicuculline-specific IRs</u>: (1) FC IR expression ≥ 20% upon bicuculline stimulation, (2) |z-score|_IR expression_ ≥ 1.5 upon bicuculline stimulation, (3) FC IR expression ≤ 5% upon BDNF stimulation; <u>BDNF-specific IRs</u>: (1) FC IR expression ≥ 20% upon BDNF stimulation, (2) |z-score|_IR expression_ ≥ 1.5 upon BDNF stimulation, (3) FC IR expression ≤ 5% upon bicuculline stimulation. For regulated intron-containing transcripts that do not belong to one of these groups, we considered we could not confidently determine their stimulation-specificity.

To probe by which RNA process IR-transcripts are regulated, we used the following criteria for the spliced isoform: <u>splicing</u>: (1) fold change mean of spliced isoform expression (FC SI expression) ≥ 20%, (2) |z-score|_SI expression_ ≥ 1.5 (where z-score_SI expression_ = (var(SI expression_unstimulated_) + var(SI expression_stimulated_)) – covar(SI expression)), (3) SI regulated in opposite directions compared to IR-isoforms; <u>degradation/stabilization</u>: (1) FC SI expression ≤ 5%, (2) |z-score|_SI expression_ ≤ 1. For regulated intron-containing transcripts that do not belong to one of these categories, we considered we could not confidently determine their type of regulation.

## Acknowledgements

We thank members of the Scheiffele lab for advice and constructive discussions. We thank Eric Allemand, Özgür Genç, Raul Ortiz and Madalena Pinto for constructive discussions and comments on the manuscript. We are grateful to Caroline Bornmann, Laetitia Burklé and Sabrina Innocenti for technical support. We thank the Quantitative Genomics Facility of Basel, in particular Philippe Demougin and Christian Beisel. We also thank Pierre De La Grange and Noémie Robil from Genosplice for support in data analysis. O.M. was financed with a Ambizione grant evaluated by the Swiss National Science Foundation. This work was supported by funds to O.M. from the Swiss National Science Foundation and to P.S. from a European Research Council Advanced Grant (SPLICECODE).

## Author contributions

M.M. and O.M. conducted the experiments and performed the computational analysis. O.M., M.M. and P.S. designed the experiments and wrote the paper.

