## Supplementary material for "Cue-specific remodeling of the neuronal transcriptome through intron retention programs": Figure Supplements and legends

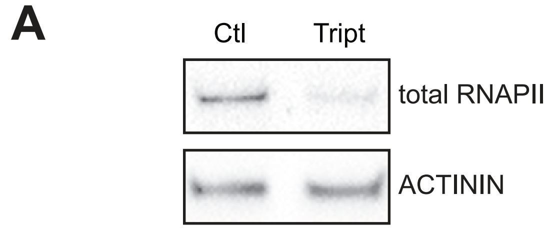


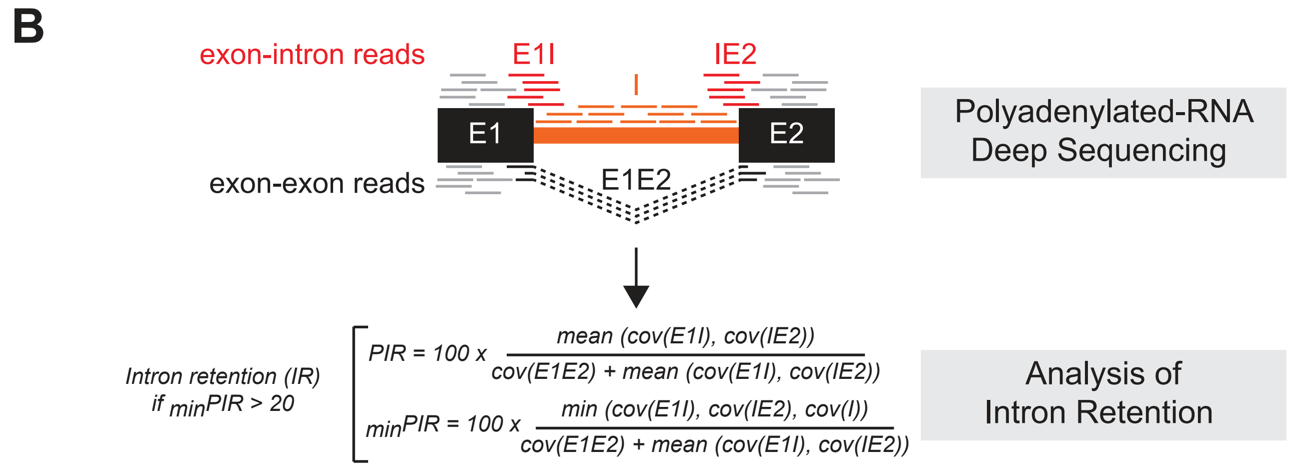


Figure supplement 1

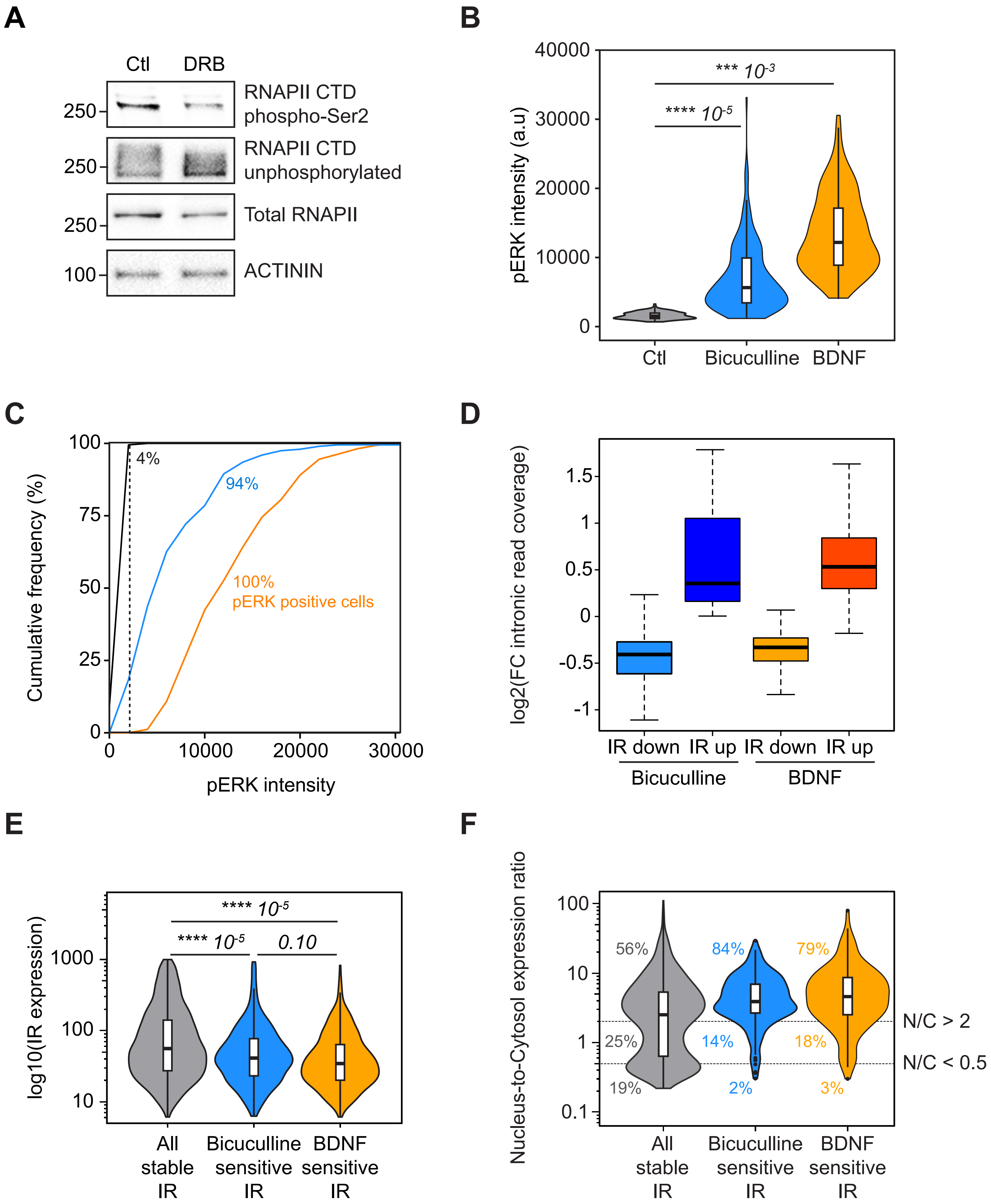


Figure supplement 2

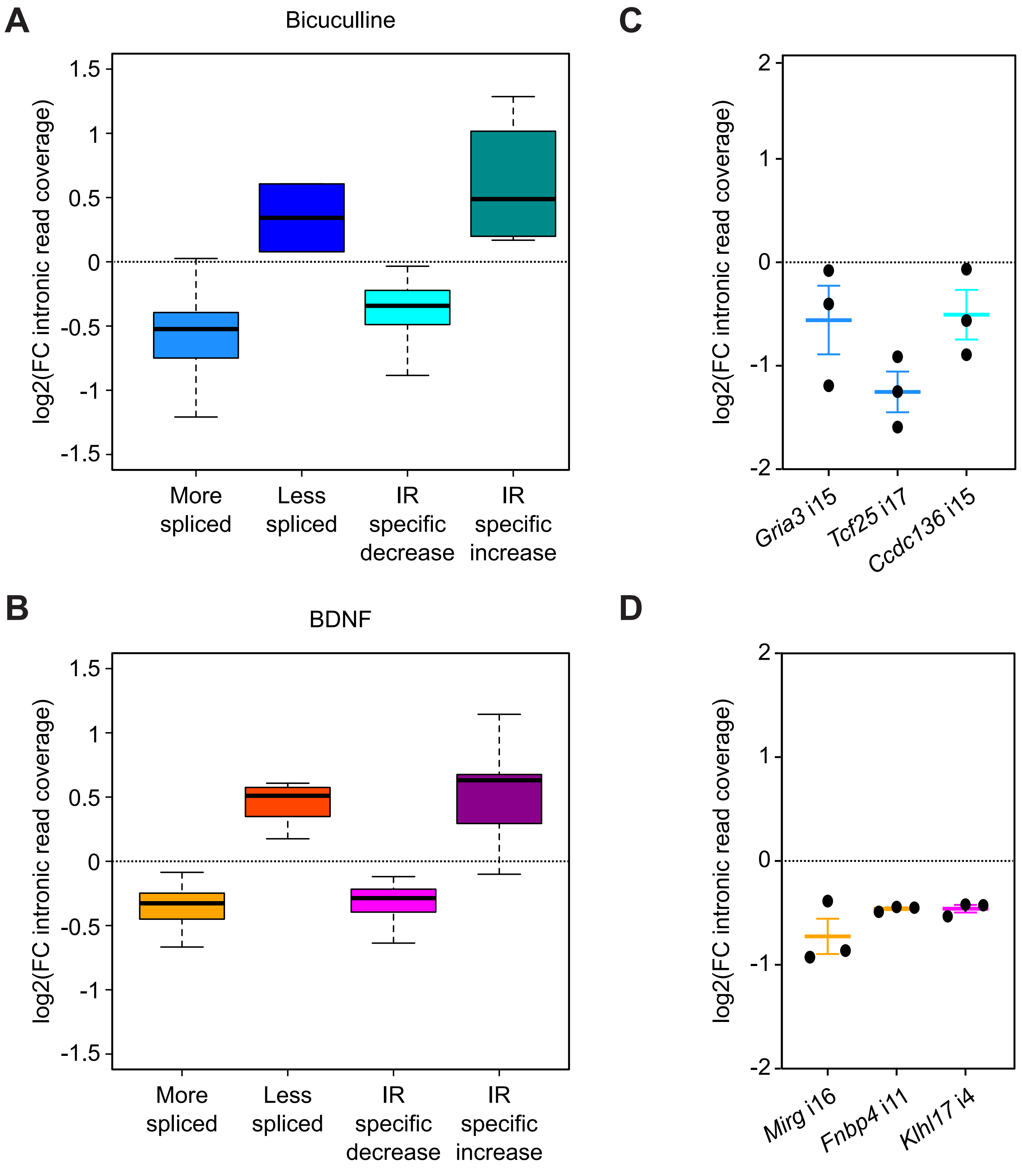


Figure supplement 3

A

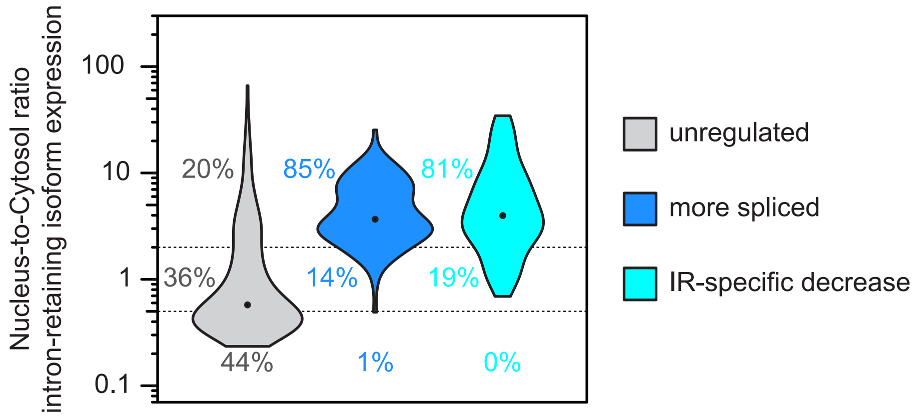


B

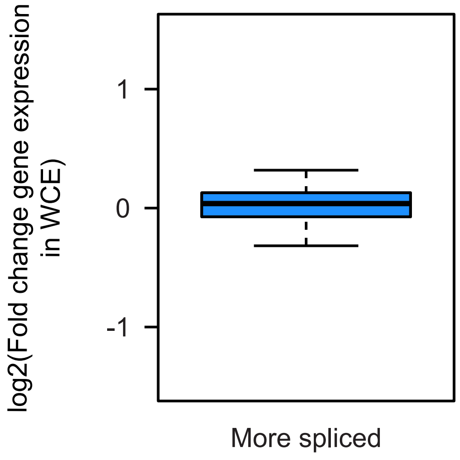


Figure supplement 4

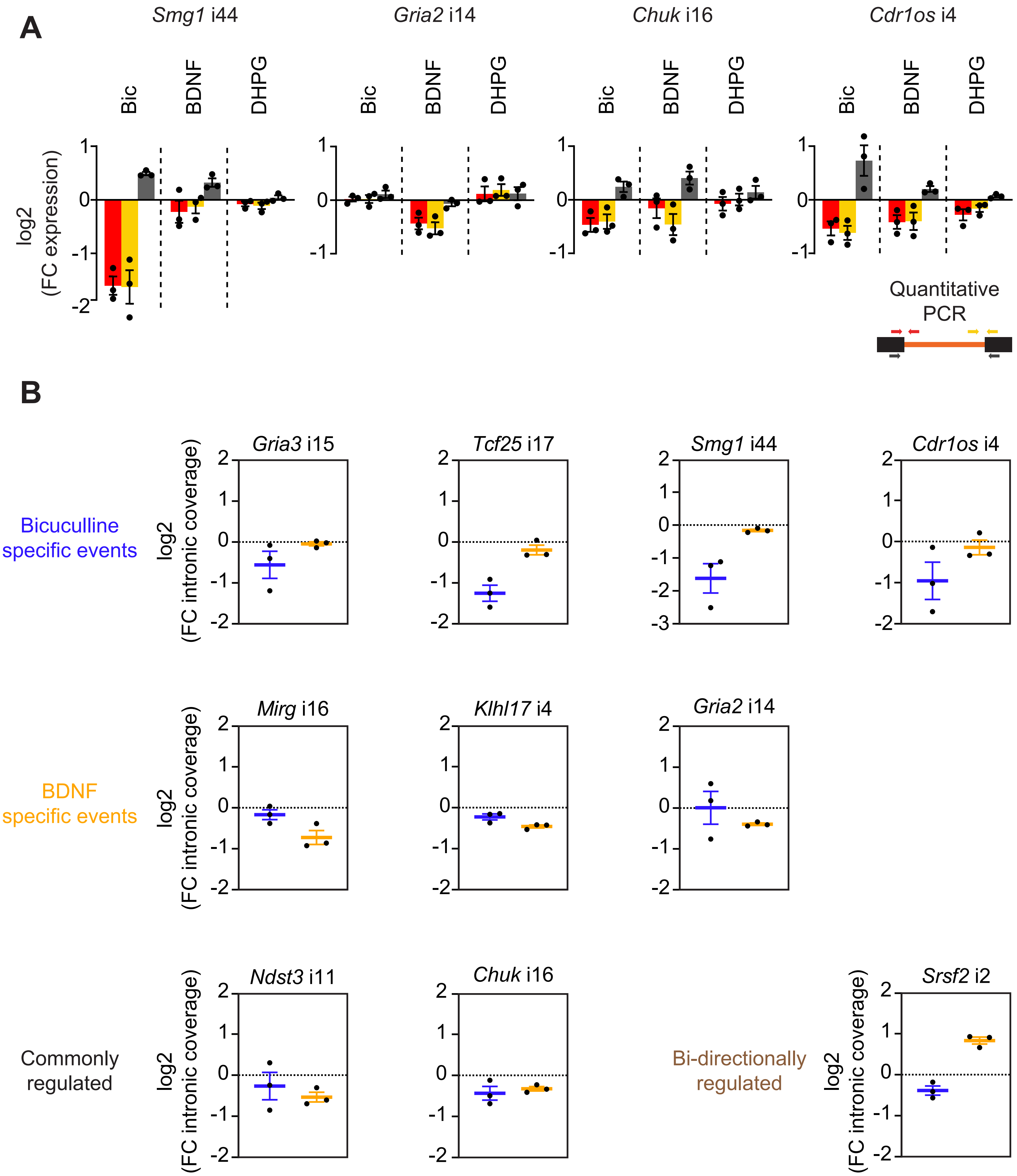


Figure supplement 5

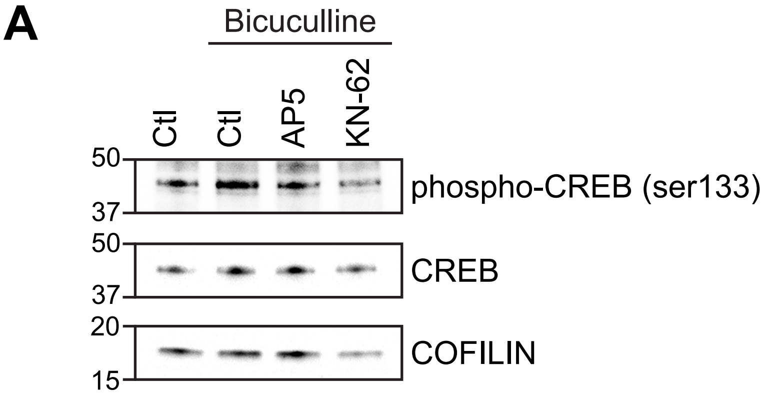


Figure supplement 6

#### Table 1 - Summary of RNA sequencing read alignment

| | | Total Reads | Uniquely mapped reads | Mapped to too many loci | Mapped to multiple loci | Unmapped reads: too short | Unmapped reads: other | RIBOSOMAL | CODING | UTR | INTRONIC | INTERGENIC | mRNA |
|---|---|---|---|---|---|---|---|---|---|---|---|---|---|
| pooled samples during sequencing | Cytosol_Ctl_Rep1 | 102381005 | 87.31% | 0.19% | 7.40% | 5.04% | 0.07% | 0.9% | 41.0% | 37.2% | 3.3% | 17.8% | 78.1% |
| | Cytosol_Ctl_Rep2 | 99258948 | 87.81% | 0.19% | 7.25% | 4.68% | 0.07% | 0.8% | 43.6% | 35.4% | 3.1% | 17.1% | 79.0% |
| | Cytosol_Ctl_Rep3 | 101852925 | 86.03% | 0.17% | 8.02% | 5.71% | 0.07% | 0.8% | 41.4% | 35.4% | 3.1% | 19.4% | 76.8% |
| | Nucleus_Ctl_Rep1 | 92173824 | 87.25% | 0.28% | 6.69% | 5.53% | 0.25% | 1.5% | 35.4% | 32.6% | 10.3% | 20.3% | 68.0% |
| | Nucleus_Ctl_Rep2 | 96510469 | 87.87% | 0.26% | 6.41% | 5.23% | 0.23% | 1.5% | 37.7% | 31.9% | 9.7% | 19.2% | 69.6% |
| | Nucleus_Ctl_Rep3 | 94707116 | 86.22% | 0.26% | 7.04% | 6.25% | 0.23% | 1.1% | 36.9% | 31.5% | 9.9% | 20.6% | 68.4% |
| | WCE_Ctl_Rep1 | 108853793 | 87.62% | 0.22% | 6.89% | 5.14% | 0.14% | 1.3% | 39.8% | 34.7% | 6.1% | 18.2% | 74.5% |
| | WCE_Ctl_Rep2 | 100207337 | 87.11% | 0.23% | 6.60% | 5.90% | 0.15% | 1.1% | 41.6% | 33.2% | 6.6% | 17.6% | 74.8% |
| | WCE_Ctl_Rep3 | 95864215 | 86.70% | 0.21% | 7.61% | 5.32% | 0.15% | 1.1% | 38.9% | 33.3% | 6.6% | 20.1% | 72.2% |
| | Cytosol_Bic_Rep1 | 88611820 | 87.28% | 0.20% | 7.07% | 5.37% | 0.08% | 0.8% | 42.4% | 36.2% | 3.5% | 17.1% | 78.6% |
| | Cytosol_Bic_Rep2 | 105010780 | 87.69% | 0.20% | 6.96% | 5.08% | 0.07% | 0.8% | 44.8% | 34.8% | 3.1% | 16.5% | 79.6% |
| | Cytosol_Bic_Rep3 | 96224784 | 87.79% | 0.19% | 6.95% | 4.99% | 0.08% | 0.7% | 43.6% | 35.8% | 3.2% | 16.8% | 79.4% |
| | Nucleus_Bic_Rep1 | 103873601 | 88.72% | 0.26% | 6.17% | 4.64% | 0.21% | 1.1% | 37.8% | 34.6% | 8.3% | 18.2% | 72.5% |
| | Nucleus_Bic_Rep2 | 91452042 | 87.14% | 0.27% | 5.92% | 6.44% | 0.24% | 1.2% | 38.5% | 32.1% | 9.9% | 18.4% | 70.5% |
| | Nucleus_Bic_Rep3 | 104217750 | 87.36% | 0.24% | 6.79% | 5.41% | 0.19% | 1.0% | 39.5% | 32.3% | 8.0% | 19.3% | 71.8% |
| | WCE_Bic_Rep1 | 99460363 | 87.90% | 0.23% | 6.37% | 5.37% | 0.14% | 1.0% | 41.0% | 35.3% | 5.8% | 16.9% | 76.3% |
| | WCE_Bic_Rep2 | 108118242 | 87.77% | 0.23% | 6.44% | 5.43% | 0.14% | 1.1% | 43.0% | 33.4% | 5.7% | 16.8% | 76.5% |
| | WCE_Bic_Rep3 | 85634802 | 87.53% | 0.22% | 6.87% | 5.27% | 0.12% | 0.9% | 42.9% | 33.8% | 5.1% | 17.4% | 76.6% |
| pooled samples during sequencing | Ctl_Rep1 | 120787493 | 88.77% | 0.25% | 6.30% | 4.60% | 0.09% | 0.9% | 48.5% | 31.7% | 5.0% | 13.9% | 80.2% |
| | Ctl_Rep2 | 115636180 | 89.20% | 0.23% | 6.18% | 4.29% | 0.10% | 0.7% | 47.2% | 31.8% | 6.2% | 14.1% | 79.0% |
| | Ctl_Rep3 | 121220186 | 88.04% | 0.23% | 6.71% | 4.92% | 0.09% | 0.7% | 46.1% | 32.8% | 5.6% | 14.9% | 78.9% |
| | Bic_Rep1 | 133680600 | 88.60% | 0.24% | 6.65% | 4.43% | 0.09% | 0.9% | 47.4% | 32.4% | 4.7% | 14.6% | 79.7% |
| | Bic_Rep2 | 106630731 | 89.55% | 0.23% | 5.84% | 4.26% | 0.12% | 0.6% | 47.1% | 31.4% | 7.2% | 13.8% | 78.5% |
| | Bic_Rep3 | 168099149 | 87.95% | 0.25% | 7.17% | 4.55% | 0.09% | 0.8% | 45.9% | 33.4% | 4.6% | 15.4% | 79.3% |
| | BDNF_Rep1 | 108996997 | 88.77% | 0.25% | 6.28% | 4.61% | 0.09% | 1.0% | 48.6% | 31.6% | 5.0% | 13.9% | 80.2% |
| | BDNF_Rep2 | 186854844 | 89.56% | 0.26% | 5.81% | 4.28% | 0.10% | 0.6% | 48.2% | 31.7% | 6.3% | 13.2% | 79.9% |
| | BDNF_Rep3 | 142143812 | 88.69% | 0.25% | 6.23% | 4.71% | 0.11% | 0.7% | 47.5% | 31.4% | 6.3% | 14.0% | 79.0% |

### Table 2- Primer sequences

EE = exon-exon junction
EI = 5' exon-intron junction
IE = 3' intron-exon junction

| | Gene | Intron | Analysed junction | Forward/ Reverse | Sequences |
|---|---|---|---|---|---|
| real-time quantitative PCR primers | Hprt | | EE | F | GATGAACCAGGTTATGACCTAGATTTGTTT |
| | | | | R | ATGGCCTCCCATCTCCTTCAT |
| | Gria3 | i15 | EI | F | CAGAGCTACAGAAAGAACAGCAGGAG |
| | | | | R | GACAGTGTGGTTCTACAACCTCTTCAA |
| | | | IE | F | CAGGTTGCCTGCAGTGTCTAAA |
| | | | | R | CGGAGTCCTTGGCTCCACATT |
| | | | EE | F | CAGAGCTACAGAAAGAACAGCAGGAG |
| | | | | R | CGGAGTCCTTGGCTCCACATT |
| | Tcf25 | i17 | EI | F | CACGGAAACACAATCGCCCTCTTCTT |
| | | | | R | GCCCACACCACTCACCTTAGAATAG |
| | | | IE | F | CCCACTTAGCTGTGTCTCATTGTTCAG |
| | | | | R | CTCCAGGTCGTTGAAGTGGAAGTT |
| | | | EE | F | CACGGAAACACAATCGCCCTCTTCTT |
| | | | | R | TCCAGCCTCTCCCCCTCTGTGG |
| | Mirg | i16 | EI | F | AGGTTGTCTGTGATGAGTTCGCTTTA |
| | | | | R | GGAAGCCTTAGACAGGGACAAACA |
| | | | IE | F | CCTACGTGTGGTAAAGCGGAAACA |
| | | | | R | ATAGGCAGGGTTCCTTGAACATCC |
| | | | EE | F | AGGTTGTCTGTGATGAGTTCGCTTTA |
| | | | | R | ATAGGCAGGGTTCCTTGAACATCC |
| | Fnbp4 | i11 | EI | F | GCAGAAGTGAATGAAGAACAAGATTA |
| | | | | R | AAAGAGCAAAGATTTCAAATAACGAAA |
| | | | IE | F | GTGTCTCTGTAGGAGAACAGATTT |
| | | | | R | GACTCTCATCTCTAACTTCCTTTGT |
| | | | EE | F | GCAGAAGTGAATGAAGAACAAGATTA |
| | | | | R | GACTCTCATCTCTAACTTCCTTTGT |
| | Ccdc136 | i15 | EI | F | GTGAGACCCTTCACAGGAGCTATG |
| | | | | R | ACCTTCCCTGCCTGACCAATAA |
| | | | IE | F | CCAGAGGTGTAGAGTGTAGGACAGA |
| | | | | R | CTCTTCCTGGCTCACTTGGTACAG |
| | | | EE | F | CCAGCAGCACAAGTGTGAGCTATAA |
| | | | | R | ACTGTTTCCTCAAAGTGCTCCAAGTC |
| | Klhl17 | i4 | EI | F | CATTCGAGGATTTGCAGACACACAC |
| | | | | R | CAAACTGGCATGTTACCTGCTTCAG |
| | | | IE | F | GCACATGCCCTCTGTCTGATACT |
| | | | | R | CGTTCAGGCTATCACTAGAGACCAATTC |
| | | | EE | F | CATTCGAGGATTTGCAGACACACAC |
| | | | | R | CCAATTCCAGCACCTGCTTCAG |
| | Smg1 | i44 | EI | F | GGGTGTAACTGGAGTAGAAGGTGTTT |
| | | | | R | TCAGTAAACTCAACACAAGTTTCCA |
| | | | IE | F | CATATGGAAACTTGTGTTGAGTTTACTG |
| | | | | R | TTCCATCTCTCGCTTACTCTGCTTG |
| | | | EE | F | TGTGAGCAGGTTCTCCACATCAT |
| | | | | R | TTCCATCTCTCGCTTACTCTGCTTG |
| | Chuk | i16 | EI | F | GAGCAGCGTGCCATTGATCTCTATAA |
| | | | | R | TACCCACTTTGTTCAGTTAGGGTGTTT |
| | | | IE | F | GGAACGTCCTGTTTGGAAATTGATGTG |
| | | | | R | GGTGACCAAACAGCTCCTTGAGAA |
| | | | EE | F | GAGCAGCGTGCCATTGATCTCTATAA |
| | | | | R | GGTGACCAAACAGCTCCTTGAGAA |
| | Srsf2 | i2 | EI | F | CTCCAGAAGAAGAGGGAGCAGTTTC |
| | | | | R | AGCATCACTCCCAAAGCTGAGTAA |
| | | | IE | F | GCTGTTTCATGCTGTTTGAGACCTATT |
| | | | | R | CCTGGAGGATCAGCCAAATCAGTTA |
| | | | EE | F | CTGCCCGAAGATCCAAGTCCAAGTC |
| | | | | R | CAGGAGACCGCAGCATTTTCTTAGGAAG |
| | Ndst3 | i11 | EI | F | AAATCCTTTGAGGAGGTACAGTTCTTT |
| | | | | R | TGTGTGGCATGAATGTTATCTGTAGT |
| | | | IE | F | TCAGTCACCAGCATATAAACGTAAGGG |
| | | | | R | GGAGCGTCCTCTGAATGGAAGTAA |
| | | | EE | F | AAATCCTTTGAGGAGGTACAGTTCTTT |
| | | | | R | GGAGCGTCCTCTGAATGGAAGTAA |
| | Smg1 | i44 | EI | F | GGGTGTAACTGGAGTAGAAGGTGTTT |
| | | | | R | TCAGTAAACTCAACACAAGTTTCCA |
| | | | IE | F | CATATGGAAACTTGTGTTGAGTTTACTG |
| | | | | R | TTCCATCTCTCGCTTACTCTGCTTG |
| | | | EE | F | TGTGAGCAGGTTCTCCACATCAT |
| | | | | R | TTCCATCTCTCGCTTACTCTGCTTG |
| | Gria2 | i14 | EI | F | GGAGTCACATTCAAGACACTGTTATTTGTT |
| | | | | R | AATCTGAATCTTTGGGTAAGGTGGTAGAG |
| | | | IE | F | GCTCACCCTGTCTGACAAGTATGTT |
| | | | | R | ACTCTCCTTTGTCGTACCACCATTT |
| | | | EE | F | TTGTTGTGGATAAATGCGGTTAACC |
| | | | | R | CTCTCCTTTGTCGTACCACCATTTG |
| | Cdr1os (C230004F18Rik) | i4 | EI | F | ACATCGCTGTGGTCCATCTCTATTTAC |
| | | | | R | ACTGGGATGGAGTAAAGGGTGAAAC |
| | | | IE | F | TCTATGTTGTGCAGAGTTACCTTATTACAC |
| | | | | R | TGTATCTCTGCTGTAGGCCAGAATTG |
| | | | EE | F | ACATCGCTGTGGTCCATCTCTATTTAC |
| | | | | R | TGTATCTCTGCTGTAGGCCAGAATTG |
| semi-quantitative PCR primers | Gria3 | i15 | | F | CAGAGCTACAGAAAGAACAGCAGGAG |
| | | | | R | CGGAGTCCTTGGCTCCACATT |
| | | | | R | TTTCCCACCCTGTTCCACCAA |
| | Tcf25 | i17 | | F | CTTCCGGTCCTTGTTGCCAAATTAC |
| | | | | R | CTCCAGGTCGTTGAAGTGGAAGTT |
| | Mirg | i16 | | F | AGGTTGTCTGTGATGAGTTCGCTTTA |
| | | | | R | ATAGGCAGGGTTCCTTGAACATCC |
| | | | | R | GGAAGCCTTAGACAGGGACAAACA |
| | Fnbp4 | i11 | | F | CCTCTGGAAGCAACTACTCCTGATTAAC |
| | | | | R | TCGCCCGACTGTTCGTTTACATAG |
| | Ccdc136 | i15 | | F | GTGAGACCCTTCACAGGAGCTATG |
| | | | | R | CTCTTCCTGGCTCACTTGGTACAG |
| | Klhl17 | i4 | | F | CATTCGAGGATTTGCAGACACACAC |
| | | | | R | CGTTCAGGCTATCACTAGAGACCAATTC |
| | Hprt | EE | | F | GATGAACCAGGTTATGACCTAGATTTGTTT |
| | | | | R | ATGGCCTCCCATCTCCTTCAT |
| | Srsf2 | i2 | | F | CTCCAGAAGAAGAGGGAGCAGTTTC |
| | | | | R | CCTGGAGGATCAGCCAAATCAGTTA |
| | Ndst3 | i11 | | F | AAATCCTTTGAGGAGGTACAGTTCTTT |
| | | | | F | TTTGCACAGGAGCTAAGGTTTGATTATT |
| | | | | R | GGAGCGTCCTCTGAATGGAAGTAA |

#### FIGURE SUPPLEMENT LEGENDS

**Figure supplement 1 – related to Figure 1**

**(A)** Activity of the RNA polymerase II inhibitor triptolide (tript) treatment was probed by assessing global expression of RNA polymerase II. Triptolide results in transcription blocks as it induces a fast proteaosome-dependent degradation of the RNA-polymerase II subunit. Mouse primary neoortical cultures were treated for 2 hours with triptolide (1µM). ACTININ was used as a loading control. (four independent cultures). These controls replicate observations made in our previous work (Mauger et al., 2016). **(B)** Polyadenylated RNAs were isolated and sequenced. For each segment comprised of two consecutive exons and the intervening intron, coverage (cov) of reads spanning the 5' and 3' exon-intron junctions (E1I and IE2, respectively), the exon-exon junction (E1E2), and mapping the intron sequence (I) were calculated. Then, the percentage of intron retention (PIR) and the minimal PIR (minPIR) were evaluated as described in the formulas. Introns were considered as retained if PIR exceeded 20 and if there was a minimum (20%) sequence coverage across the entire intron.

**Figure supplement 2 – related to Figure 2**

**(A)** Activity of RNA polymerase II inhibitor DRB treatment was probed by assessing C-terminal domain (CTD) phosphorylation and global expression of RNA polymerase II. DRB inhibits the CDK9 kinase and consequently prevent phosphorylation of the RNA polymerase II CTD (C-terminal domain). ACTININ was used as a loading control. These controls replicate observations made in our previous work (Mauger et al., 2016). **(B)** Violin plots displaying quantification of phospho-ERK signal in control (grey), bicuculline-stimulated (blue) and BDNF-stimulated mouse primary neocortical cells (orange). (four independent cultures; n=200 neurons (untreated), 201 neurons (bicuculline), 165 neurons (BDNF)) **(C)** Cumulative frequency plot displaying percentage of phospho-ERK-positive (pERK) neurons control (black), bicuculline-stimulated (blue) and BDNF-stimulated (orange) cultures. **(D)** Box plots displaying the fold change of read coverage along the whole retained intron of regulated transcripts. (three independent cultures) **(E)** Violin plots displaying the IR-transcript expression associated with stable IRs (grey), bicuculline-regulated IRs (blue) and BDNF-regulated IRs (orange). **(F)** Violin plots displaying the nucleus-to-cytosol expression ratio of IR-transcripts. All stable IR-transcripts (grey), bicuculline-regulated (blue) and BDNF-regulated transcripts are displayed. Percentage of nuclear-enriched (N/C>2) and cytosolic-enriched (N/C<0.5) transcripts are indicated on the panel. (three independent cultures). The p-values calculated with a two-sided Mann-Whitney test are indicated on the top of each panel (B and E).

**Figure supplement 3 – related to Figure 3**

**(A)** and **(B)** Boxplot displaying the fold change of read coverage along the whole retained intron in the different categories of regulated IR-transcripts upon stimulation with bicuculline (A) or BDNF (B). **(C)** and **(D)** Boxplot displaying the fold change of read coverage along the whole retained intron of transcript candidates regulated upon stimulations with bicuculline (C) or BDNF (D). Displayed transcripts correspond to those assessed by PCRs in the main Figure 3. Means and SEMs are displayed.

**Figure supplement 4 – related to Figure 4**

**(A)** Violin plots displaying the nucleus-to-cytosol expression ratio of IR-transcripts unregulated (grey), regulated through splicing (dark blue) or regulated through degradation (light blue). Percentage of nuclear-enriched (N/C>2) and cytosolic-enriched (N/C<0.5) transcripts are indicated on the panel. **(B)** Box plot displaying the fold change in the overall gene expression of gene containing a regulated retained intron upon bicuculline treatment.

**Figure supplement 5 – related to Figure 5**

**(A)** RT-PCR validations of IR-transcripts specifically regulated upon bicuculline stimulation (*Smg1*), BDNF stimulation (*Gria2*), commonly regulated (*Chuk* and *Cdr1os*). Fold changes in the expression of the IR- (red and orange) and spliced (dark grey) isoforms were assessed with real-time qPCR using three different primer sets, as represented in the bottom right scheme. SEMs are displayed (four independent cultures). **(B)** Boxplot displaying the fold change of read coverage along the whole retained intron of transcript candidates regulated upon stimulations with bicuculline and BDNF. Displayed transcripts correspond to those assessed by PCR in the main Figure 5 and figure supplement 5A. Means and SEMs are displayed.

**Figure supplement 6 – related to Figure 6**

**(A)** Efficiency of the pharmacological compounds AP5 blocking NMDA receptors (NMDAR) and KN-62 targeting the $Ca^{2+}$/calmodulin-dependent protein kinases (CaMK) was controlled by assessing level of CREB serine 133 phosphorylation by western blot; COFILIN was used as a loading control. In all conditions, DRB (50μM) was applied to mouse primary neocortical cells (14 days in culture) for 2hrs; 1hr before cell collection, bicuculline (20μM) was added to the cultures. 15min before bicuculline stimulation, the NMDAR antagonist AP5 (50μM) or CaMK blocker KN-62 (10μM) were applied. (three independent cultures) **(B)** Efficiency of the pharmacological blocker U0126 of the mitogen-activated protein kinases (MAPK) was controlled by assessing level of ERK phosphorylation by western blot; COFILIN was used as a loading control. In all conditions, DRB (50μM) was applied to mouse primary neocortical cells (14 days in culture) for 2hrs; 1hr before cell collection, BDNF (50ng/mL) was added to the cultures. 15min before BDNF stimulation, the MAPK inhibitor U0126 (10μM) was applied. (three independent cultures)

**Table 1. Summary of RNA sequencing read alignment**

**Table 2. Primer sequences**
